# Pharmacological potentiation of Nav1.1 channels in interneurons mitigates tau depositions and neuronal death in a mouse model of neurodegenerative dementias

**DOI:** 10.1101/2025.08.20.671400

**Authors:** Kazuaki Sampei, Junya Hirokawa, Takehiro Kudo, Chie Seki, Hiroyuki Takuwa, Jun Maeda, Maiko Ono, Masaki Tokunaga, Nobuhiro Nitta, Shigeyuki Honda, Taeko Kimura, Yumi Matsushita, Taizo Ishikawa, Yuhei Takado, Ming-Rong Zhang, Takafumi Minamimoto, Naruhiko Sahara, Makoto Higuchi, Masafumi Shimojo

## Abstract

Epileptiform discharges and neuronal hyperexcitability are key pathophysiological features of Alzheimer’s disease and related tauopathies. We previously identified selective dynfuntion of parvalbumin-positive GABAergic interneurons (PV neurons), which regulate neural network excitability, in a tauopathy mouse model. However, the mechanistic link between PV neuron deficits, tau pathology, and neurodegeneration remains unclear. Here, we demonstrate that pharmacological enhancement of phasic PV neuron activity markedly attenuates tau accumulation and neuronal loss in a tauopathy mouse model. We developed DSR-143630, a novel activator of voltage-gated sodium channel Nav1.1, selectively expressed in PV neurons. Administration of DSR-143630 alleviated febrile seizures in Nav1.1 haploinsufficient mice and suppressed high-frequency oscillations (HFOs), an electrophysiological signature of hyperexcitability associated with cognitive impairments, in rTg4510 tau transgenic mice. Longitudinal tau PET and volumetric MRI demonstrated that DSR-143630 treatment from 4 to 11 months of age profoundly reduced age-dependent tau deposition and atrophy in the neocortex and hippocampus. Postmortem analyses further revealed decreased levels of phosphorylated tau, preservation of neuronal populations, and attenuated neuroinflammatory responses, including reactive gliosis. These findings establish PV neuron dysfunctions and consequent network hyperexcitability as key drivers of tau pathogenesis and highlight pharmacological Nav1.1 activation as a promising disease-modifying strategy for neurodegenerative tauopathies.

**One Sentence Summary:** Pharmacological potentiation of interneuronal activities attenuates aberrant network excitability, progressive tau deposition, and neurodegeneration.

## INTRODUCTION

Progressive neuronal loss associated with the formation of senile plaques (SPs) and neurofibrillary tangles (NFTs) is a common pathological feature of Alzheimer’s disease (AD) (*1*). SPs are extracellular depositions consisting of hydrophobic amyloid-β (Aβ) peptide derived from sequential proteolysis of amyloid precursor protein (APP), while the major component of NFTs is composed of intracellular inclusions of hyperphosphorylated microtubule-associated protein tau, which are more broadly observed in diverse tauopathy spectrum disorders such as frontotemporal lobar degeneration (*2*). There has been cumulative evidence supporting the mechanistic implication of tau depositions in these illnesses, and neuropathological studies of post-mortem human brains have indicated that the NFT stage is closely correlated with the severity of synaptic degeneration and neuronal death (*3, 4*). This notion is currently supported by positron emission tomography (PET) of living tauopathy cases, demonstrating pivotal associations of tau pathologies with local brain atrophies and severity of cognitive impairments (*5, 6*). These insights into tau-provoked neurotoxicities have also raised the possibility of ameliorating neuronal viability and brain functions by therapeutic modification of tau abnormalities.

Growing evidence has indicated that the aberrant excitation-inhibition (E/I) balance of the cerebrocortical circuit underlying brain dysfunction is bidirectionally and mechanistically linked to tau accumulation at an early stage of tauopathies (*7*). A high incidence of epileptic seizures has been reported in AD and several other tauopathies (*8, 9*), and asymptomatic epileptic discharge associated with subsequent cognitive decline is frequently observed in the temporal lobe of AD patients (*10*). Likewise, independent strains of tau transgenic mice exhibit enhanced susceptibility to drug- and kindling-induced epileptic seizures at a young age, indicating that early-stage tau pathologies and related abnormalities strongly impact cortical network excitability (*11–13*). Conversely, other studies have also demonstrated that excess neuronal excitation accelerates the propagation of tau fibrillogenesis and consequent neurodegenerative processes in animal models (*14, 15*). These findings suggest that the reciprocal enhancements of tau depositions and neural network hyperexcitability may be a critical driving force in the pathogenetic events.

Notably, disturbances of the inhibitory neurotransmission mediated by γ-aminobutyric acid (GABA) are likely to be fundamentally involved in the dysregulated E/I balance in circuit activity. Although GABAergic interneurons constitute a relatively minor population of cortical neurons, they make major contributions to learning, memory formation, and various other cognitive functions via the tuning of the neuronal excitability and the maintenance of the network synchronization (*16*). The selective degeneration of GABAergic neuronal components has been documented in the brains of AD patients and model mice (*17–19*), and these abnormalities can be reversed by genetic enhancement of inhibitory neuronal activities (*19, 20*). Despite the significant roles played by the deterioration of interneurons in the neurodegenerative etiology, there has thus far been no established therapeutic approach to the modulation of the inhibitory GABAergic system in AD and allied disorders.

Nav1.1 is a subtype of the voltage-gated sodium channel (Nav) and is predominantly expressed in parvalbumin-positive GABAergic interneurons (PV neurons), which constitute a major cell population responsible for maintaining neural circuit tone and synchrony in the forebrain (*21*). Autosomal dominant loss-of-function mutations in the *Scn1a* gene encoding Nav1.1 cause Dravet syndrome, a severe myoclonic epilepsy in infancy, and genetic haploinsufficiency of Nav1.1 in mice also leads to febrile seizures with epileptiform activity due to GABAergic dysfunction (*21, 22*). Importantly, the electrophysiological and phenotypic abnormalities in a mouse model of Dravet syndrome can be significantly suppressed by Nav1.1-activating spider toxin or genetic overexpression of the multifunctional Nav β1 auxiliary subunit (*23, 24*). Moreover, we recently reported that PV neurons and/or fast-spiking interneurons were functionally disturbed along with neuroinflammation in a tauopathy mouse model (*25*). These findings provide a rationale for large-scale screening of pharmacological compounds acting on Nav family proteins as potential therapeutic agents counteracting neurodegenerative tau pathogenesis, resulting in the discovery of a candidate Nav1.1 activator dubbed DSR-143630. In this study, we demonstrate that DSR-143630 can potentiate Nav1.1 *in vivo* and mitigate tau depositions, neuronal loss, aberrant brain oscillation, and behavioral abnormalities in rTg4510 tau transgenic mice recapitulating tauopathies.

## RESULTS

### Subhead 1: DSR-143630 potentiates Nav1.1 *in vitro* and *in vivo*

DSR-143630 (**Fig. S1A**) was identified through a large-scale screen for small molecules targeting Nav family proteins, with a particular focus on Nav1.1 modulation. We first examined whether DSR-143630 potentiates Nav1.1 channel activity using whole-cell patch-clamp recording of voltage-dependent Na^+^ currents in HEK293 cells overexpressing various α subunits of human Nav channel proteins. A single-peak Na^+^ current was evoked by the voltage ramp from -100 mV to +85 mV, and the bath application of DSR-143630 efficiently enhanced the peak amplitude and prolonged the decay of its current (**Fig. 1A**). DSR-143630 most prominently potentiated Nav1.1-mediated inward currents at high concentrations, whereas it also enhanced the other Nav channels in a concentration-dependent manner (**Fig. S1B**).

**Fig. 1.**
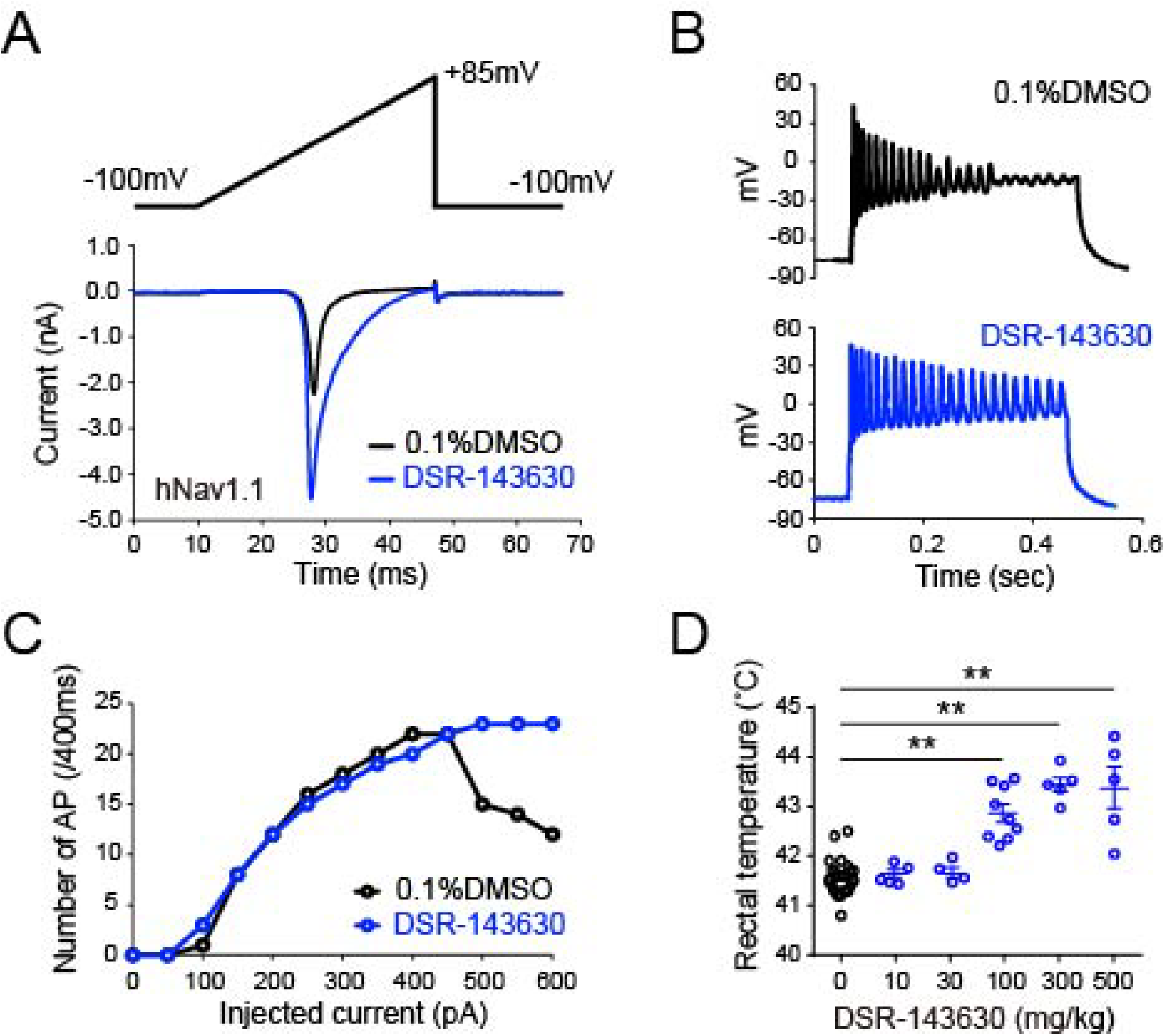
DSR-143630 potentiates Nav1.1 channel and rescues epileptiform abnormality in a mouse model of Dravet syndrome. **(A)** The pharmacological action of DSR-143630 on Nav channel was assessed by auto-patch-clamp system in HEK293 overexpressing human Nav1.1. Representative traces of Na current in the presence (blue) or absence (black) of 10 μM DSR-143630 are presented as a superimposed image. **(B-C)** Action potentials elicited by positive current injections were recorded from non-pyramidal interneurons in cortical slices of Scn1a(+/−) mice. Bath applications of 30 μM DSR-143630 (blue) or 0.1% DMSO (black) were started 10 min before the recording. **(B)** Representative traces of whole-cell, current-clamp mode recordings with current injection at 600 pA. Note that Scn1a(+/−) slices show a typical collapse of firing frequency at >500 pA (upper) and that DSR-143630 pre-treatment rescues this abnormality (lower). **(C)** The input-output curve of action potential frequency response to current injection. The number of action potentials represents the number of times the membrane potential exceeded 0 mV. **(D)** Febrile seizure phenotype observed in Scn1a(+/−) mice receiving various concentrations of DSR-143630 i.p. injection. The graph shows the rectal temperature of mice at the onset of febrile seizures. Data from the indicated number of animals in each condition are presented as Mean ± S.E.M. **p < 0.01 vs. vehicle; parametric Dunnett’s multiple comparison test.

To assess functional relevance in GABAergic interneurons, we next conducted whole-cell patch-clamp recording from non-pyramidal neurons (i.e., putative GABAergic interneurons) in acute cortical slices from Scn1a(+/−) mice, which carry a heterozygous loss-of-function mutation in Nav1.1 and serve as a model for Dravet syndrome. As reported previously (*21, 22*), stepwise current injection elicited a current-dependent increase in the frequency of action potentials at < 500 pA but caused a sudden collapse at higher conditions (**Fig. 1B and 1C**). Bath application of 30 μM DSR-143630 restored sustained firing without collapse (**Fig. 1B and 1C**), indicating that this compound rescues the channel activity of Nav1.1 to a functionally competent level in GABAergic interneurons.

We then assessed *in vivo* pharmacological effects of DSR-143630 in Scn1a(+/-) mice. Following intraperitoneal administration (100 mg/kg), brain concentration reached 18.8 μM at 60 min (**Table S1**), exceeding its *in vitro* EC50 on Nav1.1 (**Fig. S1B**) and confirming the blood-brain barrier penetration at pharmacologically effective levels. Subsequently, a heat chamber device to gradually increase the body temperature of animals was custom-built, and Scn1a(+/-) mice constantly exhibited febrile seizures under this experimental condition (**Fig. 1D**). Remarkably, intraperitoneal pre-administration of DSR-143630 efficiently suppressed the susceptibility of mice to febrile seizures in a dose-dependent manner: ≥ 100 mg/kg significantly increased the seizure threshold temperature and prolonged latency to onset (**Fig. 1D and Fig. S2**). These results demonstrate that DSR-143630 achieves effective brain exposure and suppresses cortical hyperexcitability *in vivo* via activation of Nav1.1.

### Subhead 2: DSR-143630 rescues the impairment of GABAergic interneurons in rTg4510

We previously reported that cortical GABAergic dysfunction emerges at early stages of tau pathology in rTg4510 tau transgenic mice, contributing to neurodegeneration (*18, 25*). Consistent with this, whole-cell recordings from interneurons in acute cortical slices of young rTg4510 mice (7 - 8 weeks) revealed impaired firing: action potentials failed to increase beyond 300 pA current injections (**Fig. S3**), resembling deficits observed in Dravet mice. This impairment was significantly mitigated by the bath application of 30 μM DSR-143630 (**Fig. S3**), providing a rationale for the use of this agent for the functional recovery of interneurons towards interventions targeting tau pathogenesis in rTg4510 mice. This concentration was sufficient for functional recovery of interneurons in both Dravet and rTg4510 models, and was well above the concentration in the brain that yielded anti-epileptic effects in Dravet mice. Pharmacokinetic analysis further showed that oral administration of DSR-143630 at 5.8 mg/g food yielded brain concentration above 30 µM, supporting dietary treatment as a feasible approach for subsequent experiments.

### Subhead 3: DSR-143630 restores aberrant high-frequency oscillation in the brains of rTg4510

To examine how tau pathology affects cortical network excitability, we performed in vivo epidural electroencephalography (EEG) recordings in 4-to 5-month-old head-fixed rTg4510 mice and their control littermates a self-initiated wheel running task (**Fig. 2A**). This paradigm enabled us to assess goal-directed locomotion, control the animals’ apparent behavioral state, and probe context-dependent visual cortical activity (*26*). We found that rTg4510 mice ran significantly farther beyond the required distance to obtain the reward than controls (t(193) = –2.49, p = 0.014; nonTg: 13 mice, 110 trials; rTg4510: 12 mice, 88 trials). In contrast, there were no significant differences in average running velocity or latency to initiate locomotion (**Fig. 2B**). These results indicate impaired regulation of goal-directed stopping behavior in rTg4510 mice, despite comparable overt behaviors during running and stopping periods.

**Fig. 2.**
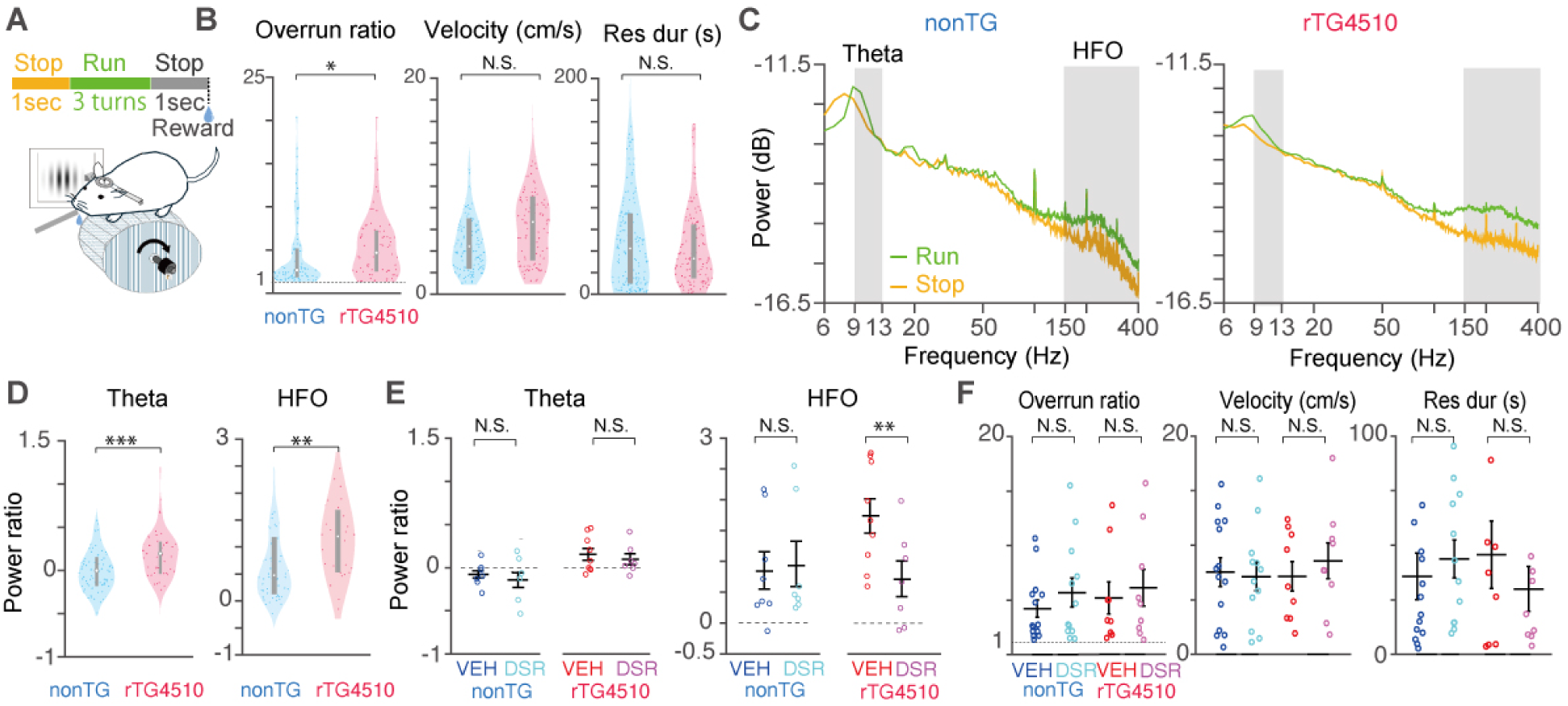
DSR-143630 alleviates the aberrant brain oscillation of rTg4510 mice. (A) Schematic of the self-initiated wheel running task. Each trial consisted of a 1-second stop, followed by ≥3 wheel turns guided by the display’s grating orientation, another 1-second stop, and a water reward delivery. (B) Overrun was defined as the ratio of actual running distance to the minimum required distance for reward delivery. rTg4510 mice at 4 – 5 months of age showed greater overrun, while running velocity and resumption duration were comparable to the controls. (C) EEG power spectra during running and stopping in example mice from control and rTg4510. Both genotypes showed enhanced theta (9–13 Hz) and high-frequency oscillation (HFO; 150–400 Hz) power during running. (D) Quantification of run/stop power ratios reveals elevated theta and HFO power in rTg4510 mice relative to the controls. (E) A single intraperitoneal injection of DSR-143630 reduced run/stop power ratios of HFO in rTg4510 mice without affecting theta. (F) DSR-143630 administration had no significant effect on behavioral performance. Data are presented as mean ± SEM. Each dot represents one animal.

We then examined EEG spectral dynamics across behavioral states in the recorded visual cortical areas. Power spectral density (PSD) plots from control and rTg4510 mice showed distinct state-dependent changes, with noticeable bumps in the theta (9–13 Hz) and high-frequency oscillation (HFO, 150–400 Hz) ranges during self-initiated running compared to stopping periods, which were especially prominent in rTg4510 mice (**Fig. 2C**). Quantitative analysis confirmed that rTg4510 mice exhibited significantly higher power in both the theta (t(125) = –3.51, p = 6.1 × 10⁻⁴) and HFO (t(125) = –2.95, p = 0.0038) bands during running relative to control littermates (nonTG:13mice, 78trials, rTG:9mice, 49trials, **Fig. 2D**).

To assess potential confounds from movement-related signals on the observed HFOs, we quantified their temporal correlation with EMG signals recorded from the trapezius muscle and found no significant correlation (**Fig. S4**). In addition, the detected HFOs exhibited multi-cycle oscillatory waveforms with a stereotyped structure that aligned with single-unit spiking activity, and were reliably observed on a trial-by-trial basis during task performance, particularly during running, suggesting structured neural activity reflecting underlying network hyperexcitability (**Fig. S4**).

Following intraperitoneal injection of DSR-143630 (30 mg/kg), HFO power in head-fixed rTg4510 mice was significantly reduced from 30 minutes post-injection, as assessed by a linear mixed-effects model accounting for repeated measurements from the same animals (nonTG: 4 mice, 12 sessions; rTg4510: 6 mice, 18 sessions). The model revealed a significant Genotype × Injection interaction (p = 0.015), indicating that the treatment selectively decreased HFO power in rTg4510 mice to levels comparable to control mice (Fig. 2E), supporting normalization of aberrant high-frequency activity by potentiation of Nav1.1. In contrast, theta power remained unaffected by DSR-143630 administration with no significant main effect of Injection (p = 0.31) nor a Genotype × Injection interaction (p = 0.94, **Fig. 2E**). Since aberrant HFOs observed in the brains of rTg4510 mice were potentially caused by dysfunction of PV neurons which predominantly express Nav1.1 channel, we examined EEG spectrum during the inactivation of cortical PV neurons by inhibitory Designer Receptors Exclusively Activated by Designer Drugs (hM4Di-DREADD). Indeed, chemogenetic suppression of PV neurons led to an increase in HFOs (**Fig. S5**), establishing a causal link between impaired PV neuron activity, Nav1.1 dysfunction, and network hyperexcitability. Notably, DSR-143630 administration did not significantly alter behavioral measures of task engagement (**Fig. 2F**), confirming that EEG changes were not secondary to overt behavioral modifications.

### Subhead 4: Long-term treatment with DSR-143630 suppresses tau depositions and brain atrophy in rTg4510 mice

We therefore carried out long-term oral administration of DSR-143630 to rTg4510 mice from 4 months of age and longitudinally monitored tau pathology and brain atrophy using [^18^F]PM-PBB3 PET (*6*) and volumetric MRI, respectively. PET imaging showed that radioactive signals in the rTg4510 brains were low at 4 months of age (**Fig. 3A**), comparable to those of the age-matched non-Tg controls, but progressively increased with age, showing 39.4% and 36.1% higher cortical and hippocampal SUVRs at 8 months (**Fig. 3A-C**). Notably, the treatment of rTg4510 mice with DSR-143630 suppressed the accumulation of tau-related radioactive signals in these brain areas throughout the observation period until 9 – 11 months of age (**Fig. 3A**). The rate of the SUVR increment in the cortex and hippocampus was declined by approximately 50% in the DSR-143630-treated groups (**Fig. 3B and 3C**), leading to reduction of the cortical and hippocampal radioligand binding estimated as (SUVR – 1) by 55.5% and 47.3% at 8 months of age.

**Fig. 3.**
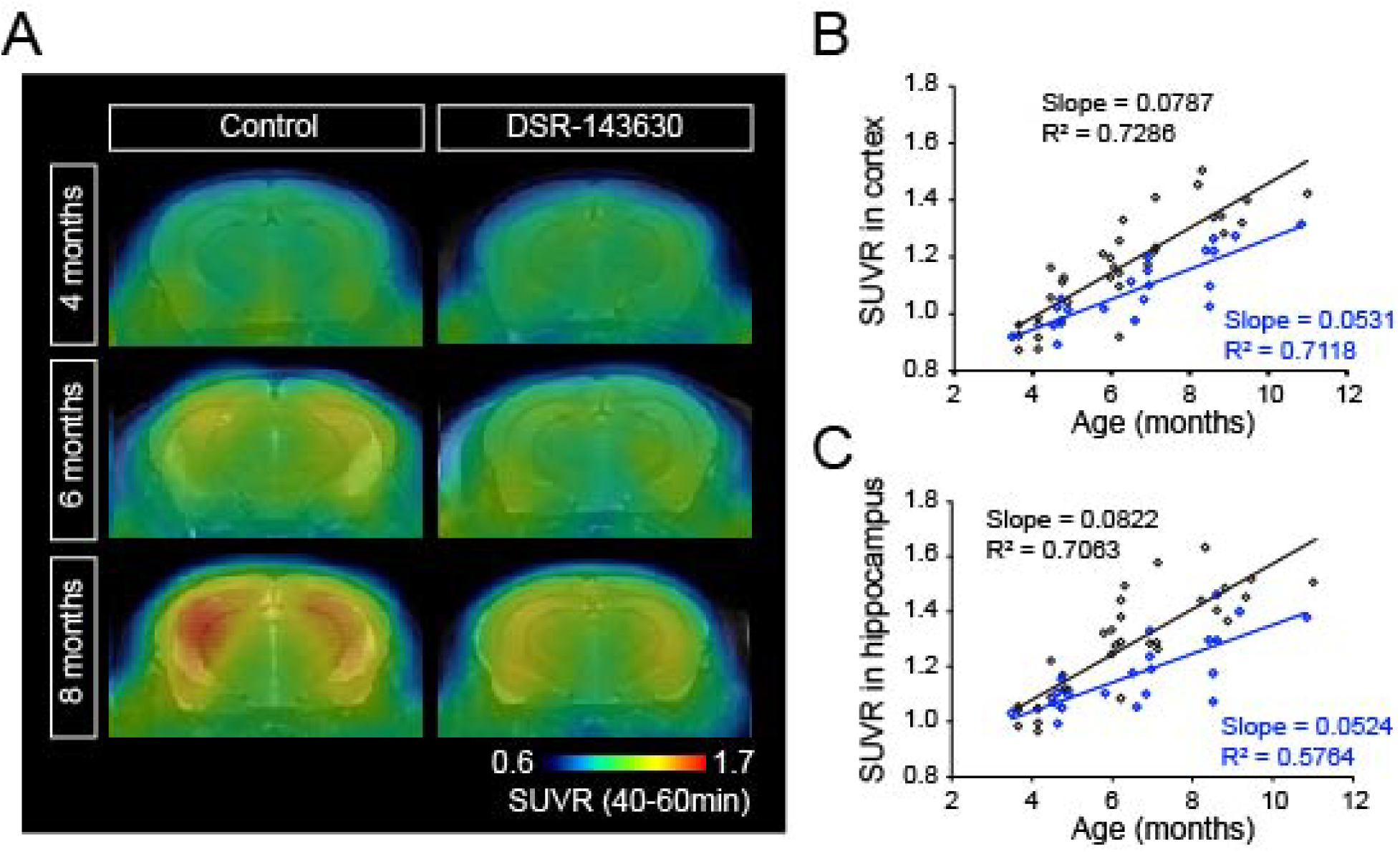
Longitudinal [^18^F]PM-PBB3 tau PET imaging of rTg4510 mice chronically treated with DSR-143630. **(A)** Representative PET images captured with [^18^F]PM-PBB3 from 4-, 6-, and 8-month-old rTg4510 mice chronically treated with a control diet or DSR-143630-containing diet. T2-weighted MR images from each group are overlaid for spatial alignment. Radioactive signal accumulation is presented as heat map images of SUVR averaged at 40-60 min after bolus injection of the tracer. **(B-C)** Progress of tau accumulation in rTg4510 mice was non-invasively assessed by PET imaging with [^18^F]PM-PBB3 during aging. Linear regression analysis shows a significant protective effect of DSR-143630 treatment in the cortex **(B)** and hippocampus **(C)** (ANCOVA, p = 0.044 for cortex, 0.045 for hippocampus). Each circle represents an individual mouse of control group (black) and DSR-143630 group (blue), respectively.

We also sequentially monitored the progressive brain atrophy of rTg4510 mice by T2-weighted MRI in parallel with PET imaging. Consistent with our previous findings (*27*), age-dependent atrophy of the cerebral cortex and hippocampus in concurrence with enlargement of the lateral ventricles was observed in control groups (**Fig. 4A**). DSR-143630 treatment significantly preserved cortical and hippocampal volumes, and the rate of age-related volume loss in both brain regions was attenuated by approximately 38% and 70%, respectively (**Fig. 4B-C**). To our surprise, the ventricular dilation was almost completely suppressed by the administration of DSR-143630 throughout aging (**Fig. 4D**). No volume change of the cerebellum was observed in either of the two groups during aging (**Fig. 4E**). Together, these results indicate that long-term potentiation of Nav1.1 by DSR-143630 mitigates tau accumulation and protects against neurodegeneration in rTg4510 mice.

**Fig. 4.**
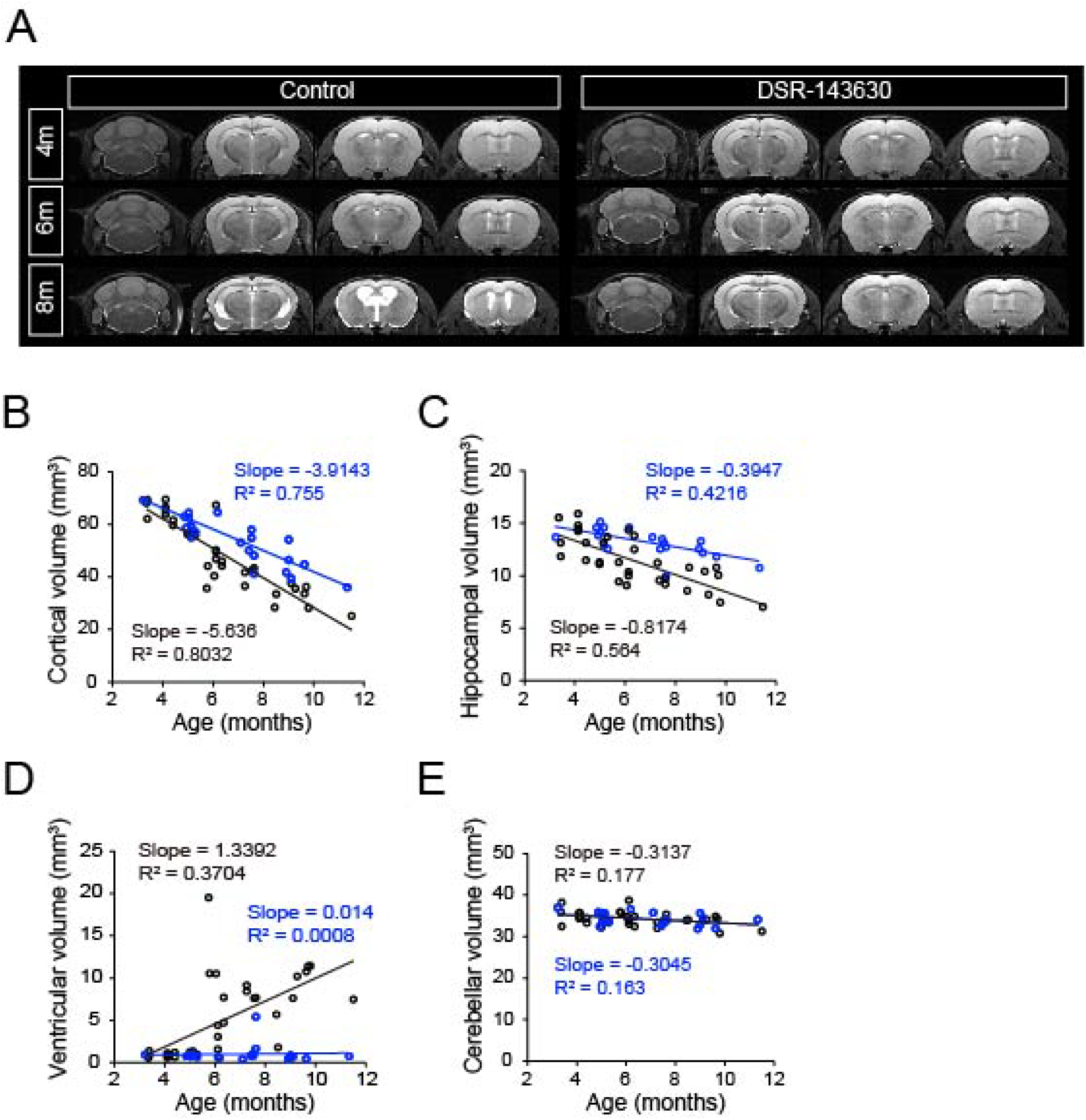
Effect of DSR-143630 on brain atrophy observed in rTg4510 mice. **(A)** Representative T2-weighted MR images of four trans-axial sections including the cerebellum, hippocampus, cortex, and striatal region of rTg4510 mice chronically treated with a control diet or DSR-143630-containing diet. **(B-C)** Progressive reduction of cortical and hippocampal volume in rTg4510 mice during aging. Linear regression analysis between the control group (black) and DSR-143630 group (blue) shows a significant protective effect of DSR-143630 treatment (ANCOVA, p = 0.0139 for cortex, 0.0156 for hippocampus). Each circle represents an individual mouse of control group (black) and DSR-143630 group (blue), respectively. **(D)** Progressive enlargement of the lateral ventricles in rTg4510 mice during aging. Note the significant protective effect of DSR-143630 treatment by linear regression analysis between two groups (ANCOVA, p = 0.0032 for lateral ventricle). **(E)** Cerebellum volume of rTg4510 brains during the experimental period.

### Subhead 5: Postmortem assessments of rTg4510 brain tissues demonstrate therapeutic blockade of tau pathologies by DSR-143630

To support our findings from PET and MRI analyses, we performed histopathological analyses of tau accumulation in post-mortem brain tissues collected from rTg4510 mice chronically fed control or DSR-143630 diet. In agreement with previous studies (*28*), immunostaining with AT8 antibody against phosphorylated tau-specific epitopes observed in diverse tauopathies demonstrated numerous NFT-like somatodendritic tau depositions in neurons, along with tau-loaded dystrophic neurites, broadly distributed to the neocortex and hippocampus of rTg4510 mice at 9-11 months of age (**Fig. 5A**). Although the number of neuronal somas harboring AT8^+^ tau deposits was unchanged, neuritic phospho-tau signals were profoundly reduced in the treatment group. Consequently, the total intensity of AT8^+^ fluorescence staining in the neocortex decreased significantly (**Fig. 5B**), and correlated with the retention of [^18^F]PM-PBB3 in PET assays at 8 months of age (**Fig. 5C**). To further examine the effect of DSR-143630 treatment on the deposition of mature tau fibrils, we also performed Gallyas-Braak silver staining. As described previously (*28*), argyrophilic neurons were abundantly present in the neocortex and hippocampus of rTg4510 mice (**Fig. 5D**). The number of Gallyas^+^ neuronal somas in the entire neocortical area was not significantly altered by the DSR-143630 treatment (**Fig. 5E**), suggesting that the compound treatment counteracts the tau fibrillogenesis in the neuritic rather than somatic compartment of neurons.

**Fig. 5.**
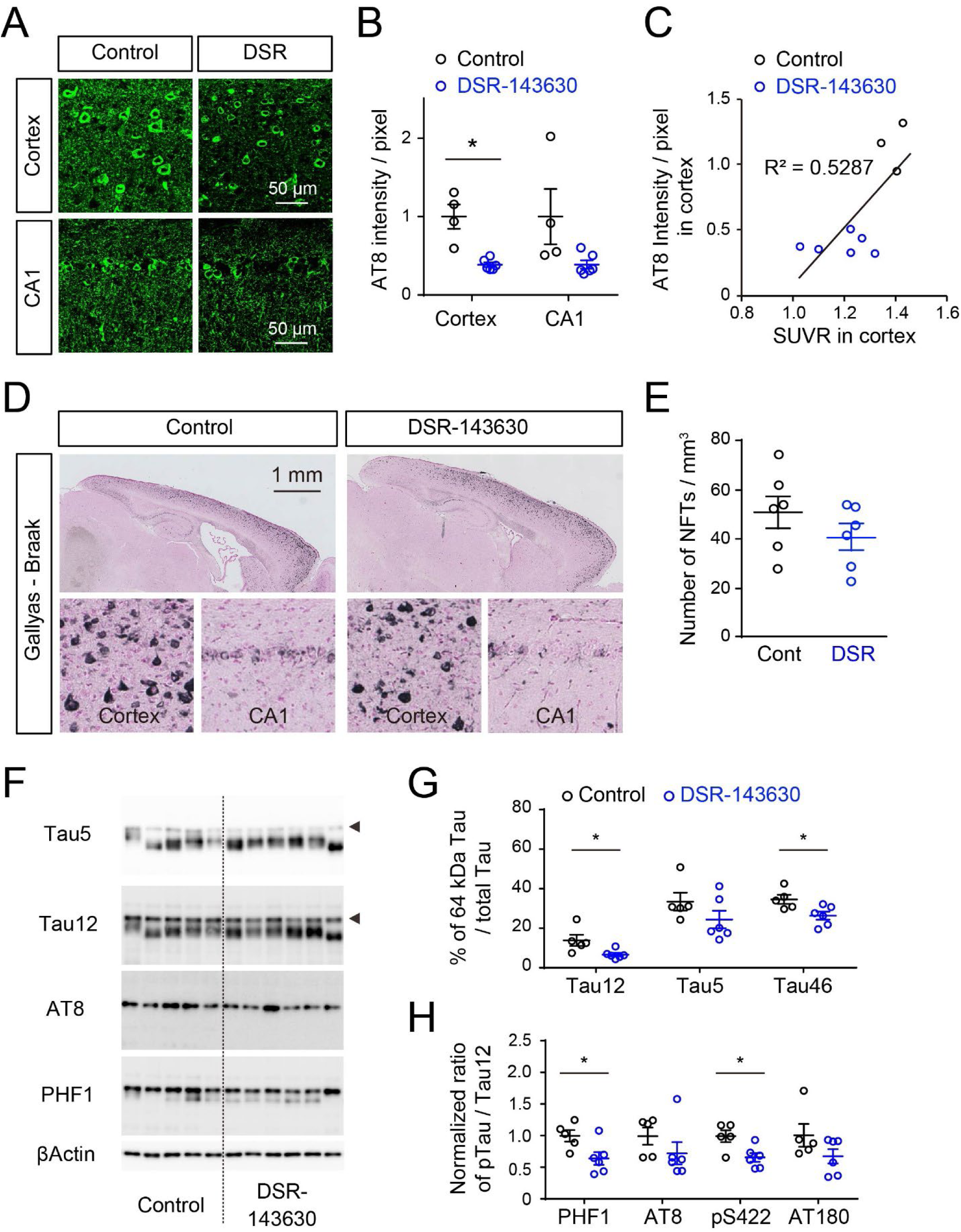
Postmortem assessment of tau pathology in rTg4510 brains chronically treated with DSR-143630 (A-C) Pathological deposition of phosphorylated tau in post-mortem brain tissues from 9-11-month-old rTg4510 mice chronically treated with control diet or DSR-143630 diet was analyzed by immunohistochemistry. **(A)** Sagittal brain section derived from one each of control diet and DSR-143630 diet rTg4510 mice immunolabeled with anti-phospho tau AT8 antibody. Representative images in neocortex (upper) and hippocampal CA1 (lower) region are presented. **(B)** Immunoreactive fluorescence intensity of AT8^+^ phosphorylated tau in the neocortex and hippocampal CA1 of rTg4510 mice. Data from control group (black, n = 4) and DSR-143630 group (blue, n = 6) are presented as Mean ± S.E.M. Each circle represents an individual mouse. Mean value of control mice was set as 1. *p < 0.05 (Student t-test) **(C)** Scatter plot for SUVR of [^18^F]PM-PBB3 PET at 8 months of age versus AT8^+^ pTau immunoreactivity in post-mortem neocortical slice among rTg4510 mice with or without DSR-143630 administration. Each circle represents an individual mouse. **(D)** Gallyas-Braak silver staining of NFT-like structures in sagittal brain sections derived from rTg4510 mice treated with 0.5% MC or DSR-143630. Representative low-magnification (upper) and high-magnification (lower) images of each condition demonstrate drug treatment suppressing the density of NFTs. **(E)** The number of NFT-like structure was quantified in the cortex. Data from the control group (black, n = 6) and DSR-143630 group (blue, n = 6) are demonstrated as Mean ± S.E.M. Each circle represents an individual mouse. **(F)** Proteins in TBS-extractable fractions were isolated from forebrain homogenates of 9-11-month-old rTg4510 mice chronically treated with a control diet and DSR-143630 diet and separated by SDS-PAGE followed by immunoblot analysis. Blots probed with various antibodies against pan-tau (Tau5 and Tau12), phosphorylated tau (AT8 and PHF1), and βactin as loading control are shown. Arrowheads indicate 64kDa tau representing hyper-phosphorylated tau species. **(G)** Relative levels of 64kDa tau normalized by levels of total tau detected by Tau5, Tau46, and Tau12 antibodies. Note that the DSR-143630 group demonstrates significantly lower levels of 64kDa tau species. Data from control group (black, n = 5) and DSR-143630 group (blue, n = 6) are plotted as Mean ± S.E.M, and each circle represents an individual mouse. *p < 0.05 (Student t-test). **(H)** Relative levels of PHF1, AT8, pS422, and AT180 positive hyper-phosphorylated tau species normalized by levels of total tau detected by Tau12 antibody. Data from control group (black, n = 5) and DSR-143630 group (blue, n = 6) are plotted as Mean ± S.E.M. Each circle represents an individual mouse. Mean value of control group was set as 1. *p < 0.05, (Student t-test).

We also extracted proteins from post-mortem brain tissues followed by immunoblotting with various tau antibodies to investigate the effect of DSR-143630 on the biochemical properties of overexpressed tau proteins. As characterized previously (*29*), human P301L tau proteins in rTg4510 mice were detected as two distinct species weighing 50-60 kDa and 64 kDa in the TBS soluble (S1) fraction, which represent physiological and hyper-phosphorylated tau species, respectively (**Fig. 5F**). Interestingly, the levels of 64 kDa tau species in the DSR-143630 group were significantly lower than those in the control group (**Fig. 5G**). According to this observation, immunoblot analysis with phosphorylation site-specific tau antibodies also demonstrated a significant reduction of multiple phospho-tau epitopes of human P301L tau in the brains of the DSR-143630-treated group relative to the controls (**Fig. 5H**). In contrast, no obvious changes in the amount of sarkosyl-insoluble tau species were observed under this experimental condition. Given these histopathological and biochemical data, it is likely that the treatment with DSR-143630 markedly inhibits the accumulation of relatively immature tau filaments in neuritic processes but may not reduce mature, insoluble tau aggregates in neuronal cell bodies due to improved survival of tau-burdened neurons.

### Subhead 6: DSR-143630 attenuates progressive gliosis and neuronal loss in rTg4510 brains

Accumulating evidence has indicated that reactive gliosis is critically involved in the progressive neuronal loss provoked by tau pathologies (*27, 30, 31*). We accordingly examined the relationship between neuronal loss and gliosis in the brains of rTg4510 mice at 6 and 9 - 11 months of age following oral DSR-143630 treatment starting at 4 months of age. Consistent with the MRI findings, cortical tissue weight and thickness were significantly preserved at 9 – 11 months by the chronic administration of DSR-143630 (**Figs. 6A-6C**). The density of NeuN^+^ neurons in the sensory cortex was also significantly preserved in treated mice compared to controls (**Fig. S6**).

**Fig. 6.**
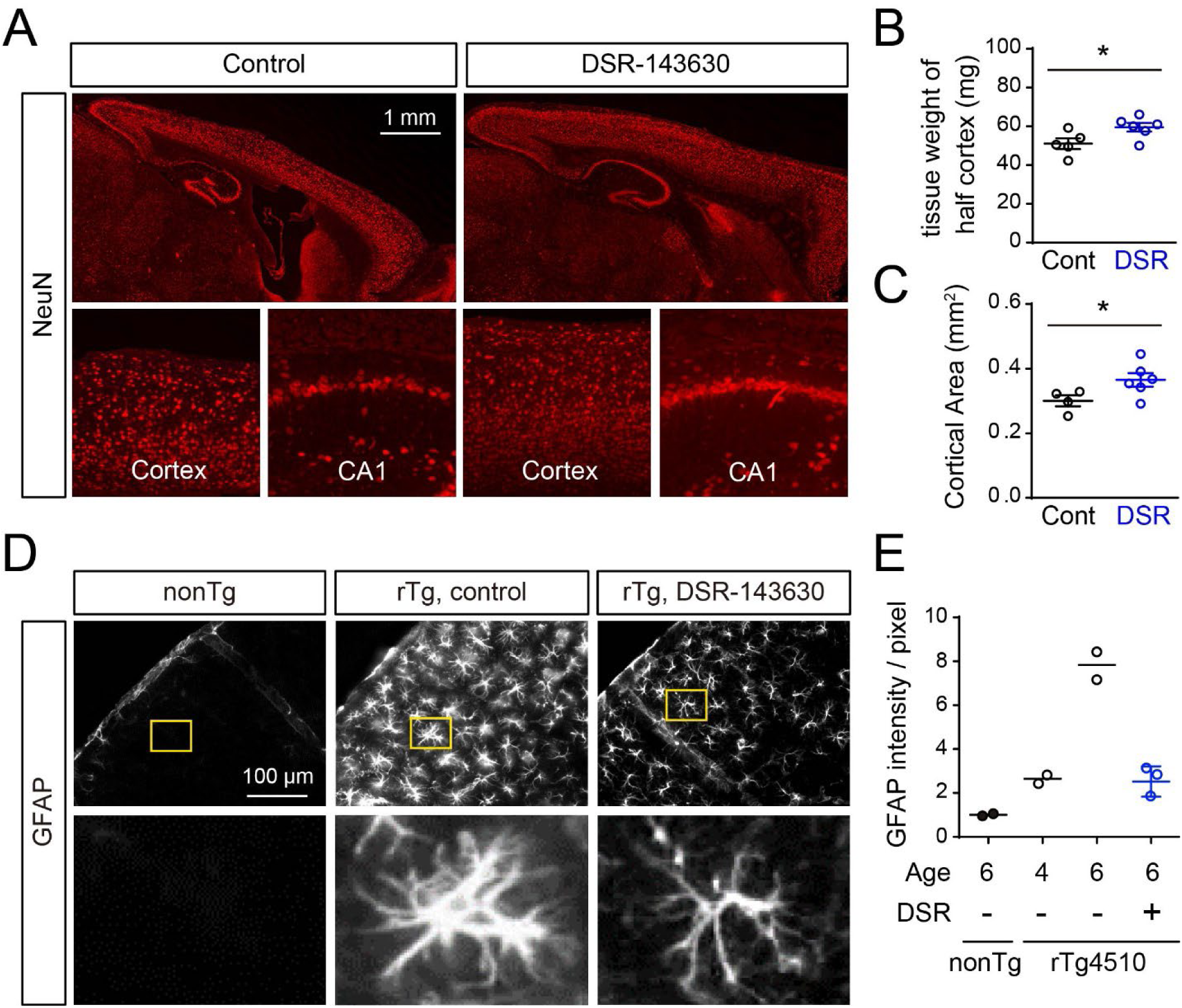
Neuronal loss and reactive gliosis in rTg4510 brains chronically treated with control diet and DSR-143630 diet (A-C) Atrophy and neuronal loss in post-mortem brain tissues from 9-11-month-old rTg4510 mice that received chronic administration of control diet (left) or DSR-143630 diet (right) were analyzed by immunohistochemistry. **(A)** Sagittal brain sections of each mouse immunolabeled with anti-NeuN antibody. Representative low- and high-magnification fluorescence images demonstrate that DSR-143630 treatment recovers progressive atrophy and neuronal loss in the neocortex and hippocampus. **(B)** Cortical tissue weight of rTg4510 mice treated with control diet (black) and DSR-143630 diet (blue). Data from 5-6 mice/genotype are demonstrated as Mean ± S.E.M. *p < 0.05, (Student t-test). **(C)** Cortical area of sagittal sections from rTg4510 mice treated with control diet and DSR-143630 diet. The area values of four brain slices from each mouse were averaged for quantification. Data from control group (black, n = 4) and DSR-143630 group (blue, n = 6) are presented as Mean ± S.E.M. *p < 0.05 (Student t-test with Welch correction). **(D)** Reactive gliosis of GFAP^+^ astrocytes in the somatosensory cortex of 6-month-old non-Tg (left) and rTg4510 mice treated with control diet (middle) or DSR-143630 diet (right) for 2 months. Representative low-magnification (upper) and high-magnification (lower) images of each condition demonstrate that drug treatment drastically suppresses the immunoreactivity. **(E)** Immunoreactive intensity of GFAP in the neocortical region of rTg4510 mice. Data from nonTg (6 months, n = 2), rTg4510 with control diet (4 months, n = 2; 6 months, n = 2), or rTg4510 with DSR-143630 diet (6 months, n = 3) are presented as Mean ± S.E.M. The Mean value of non-Tg mice was set as 1.

We then investigated the immunoreactivity of these tissue slices with GFAP and Iba1, which are representative markers of astrocytes and microglia, respectively. As reported previously (*27*), cortical GFAP signals were markedly enhanced in 6-month-old rTg4510 mice fed the control diet compared with non-Tg mice (**Fig. 6D, E**). This astrocytic activation in the cortex was significantly reduced in rTg4510 mice treated with DSR-143630 (**Fig. 6D, E**). In contrast, at 9 – 11 months of age, GFAP and Iba1 immunolabeling did not differ between treated and control groups, suggesting that DSR-143630 suppresses early gliotic response to relatively immature tau aggregates but has limited effect once high-order tau assemblies have accumulated.

### Subhead 7: DSR-143630 treatment restores the cognitive functions of rTg4510 mice

Finally, we tested whether DSR-143630 administration can restore cognitive function using a novel object recognition task. rTg4510 mice at 3.5 months were fed either control or DSR-143630 diet for 2 months, with age-matched non-Tg mice as controls. At 4.5 months, non-Tg mice displayed intact recognition memory, spending more time exploring a novel object (**Fig. S7**). Consistent with a previous study (*32*), rTg4510 mice fed the control diet failed to show novel object preference, indicating impaired short-term episodic memory. In contrast, rTg4510 mice receiving the DSR-143630 diet exhibited normal preference comparable to non-Tg mice (**Fig. S7**), indicating that the potentiation of GABAergic interneurons alleviates tau-induced cognitive deficits.

## DISCUSSION

Increased network excitability has been observed in AD patients and animal models (*11, 33*) and has been implicated as an accelerator of neuronal deterioration in AD pathogenesis (*33*). These epileptic activities of neurons are multifactorially attributable to dysfunction of GABAergic interneurons and/or degeneration in inhibitory synapses (*17, 18, 25, 34*). Meanwhile, there have been no therapeutic approaches to Aβ- and tau-triggered neurodegenerative pathways by repressing aberrant neuronal hyperexcitability without disturbing the physiological activities of excitatory neurons. Here, we utilized DSR-143630, which was capable of augmenting Nav1.1 functions in PV neurons, for the suppression of epileptiform discharges in mouse models of Dravet syndrome and to reduce high-frequency cortical network hyperactivity in rTg4510 tauopathy model mice. Notably, longitudinal neuroimaging assessments demonstrated a pronounced diminution of tau depositions and neuronal death, leading to rescue of brain functions as examined by behavioral phenotypes. Hence, the beneficial effects of the Nav1.1 activator are not confined to neuroprotection against excitotoxicity, and our strategy opens an avenue to anti-tau therapeutics.

HFOs have been extensively studied as biomarkers of epileptiform activity, particularly in the hippocampus and entorhinal cortex of epilepsy patients and rodent models (*35, 36*). However, recent studies have also highlighted the presence of physiological HFOs that are transiently induced during memory encoding and cognitive processing (*37*). In light of this, the increased HFO power observed in rTg4510 mice during self-initiated wheel running is unlikely to reflect overt epileptic discharges, given their trial-by-trial consistency, structured morphology, and independence from EMG activity (Fig. S4). Similar findings have been reported in other AD model mice such as Tg2576, which show elevated cortical HFOs without seizures (*38*). In addition, a recent clinical study in AD patients demonstrated increased HFOs even in the absence of epilepsy, suggesting that HFO augmentation does not simply reflect epileptogenesis (*39*). Instead, these changes likely arise from distinct network alterations, linked to PV neuron dysfunction (*25*) (Fig. S5).

Pharmacological suppression of hyperexcitability may offer a promising strategy to prevent or delay the onset of cognitive decline in Alzheimer’s disease. Importantly, our EEG recordings were conducted at 4–5 months of age, where overt tau aggregation and neuronal loss are not yet prominent (*28, 29*). The presence of aberrant HFO activity and cortical hyperexcitability during this pre-degenerative window suggests that network dysfunction may precede and possibly facilitate the progression of tau pathology. These findings support the emerging view that functional circuit alterations may serve as early indicators and possibly drivers of neurodegenerative processes, even before overt brain atrophy becomes apparent. Notably, while DSR-143630 improved novel object recognition, it failed to reduce overrun in rTg4510 mice (Fig. 2F). This suggests that acute intervention may be sufficient to rescue spontaneous innate behaviors, but not overtrained, habitual behaviors such as goal-directed locomotor control, which may require longer-term modulation. Taken together, these findings suggest that early pharmacological suppression of cortical hyperexcitability, possibly through modulation of PV neuron function, may help prevent or partially reverse cognitive decline associated with tau pathology, offering a potential therapeutic window before irreversible degeneration ensues.

Although the mechanisms by which DSR-143630 attenuated the accumulation of tau aggregates are yet to be unraveled, the anti-epileptic action of this potential drug may decelerate the propagation of pathological tau species in a transsynaptic fashion. The prion-like dissemination of misfolded tau molecules has been indicated in diverse neurological disorders (*40*), and investigations of mouse models receiving inoculation of tau fibrils (*41–44*) and locally expressing transgenic tau proteins (*45, 46*) have demonstrated the transfer of tau pathologies via neural circuit connections. It is noteworthy that the release of tau to the extracellular matrix was found to be reinforced by the topical application of epileptogenic agents in a previous microdialysis experiment (*47*). Likewise, propagation of tau pathologies was reported to be promoted when neurons were depolarized by high-frequency optogenetic stimulation and chemogenetic excitation (*14*), which were indicated to be relevant to epileptogenic conditions (*48, 49*). An alternative mechanism linking neuronal hyperexcitability and tau accumulation could be an axis constituted of glutamate receptor, Fyn kinase, and tau in dendritic spines (*50*), provoking calcium overflow and hyperphosphorylation of tau by Cdk5 and Fyn (*51, 52*).

In line with these notions, immunohistochemical assessments in the current work revealed a preferential reduction of neuritic versus somatic inclusions of phosphorylated tau proteins by the treatment with DSR-143630. Immunoblotting analyses also illustrated treatment-induced declines of phosphorylated tau species in the soluble protein fraction, which could consist of misfolded tau monomers and oligomers. Since an oligomeric form of tau molecules has been shown to exert neurotoxicity more pronouncedly than packaged high-order assemblies (*53–55*), an agent counteracting early-stage tau multimerization should be efficacious for the rescue of neurons from deleterious insults. It has also been documented that cell-to-cell propagation of such oligomers is a crucial process in the dissemination of neurodegenerative proteinopathy (*53, 54*), whereas it remains undetermined whether tau assemblies are transmitted at synapses in a manner dependent on the neuronal excitation. The suppression of tau depositions was also demonstrated by longitudinal PET measurements with a specific radioprobe, [^18^F]PM-PBB3. It was somewhat unexpected that the probe retention was decreased by approximately 50% at each age despite no overt reduction of sarkosyl-insoluble tau aggregates as examined by immunoblots. As the access of the probe to the inner portion of a bulky tau deposit in the neuronal soma could be impeded in the living brain, tau PET imaging may be sensitive to changes in neuritic rather than somatic tau inclusions. By contrast, large-sized somatic tau fibrils could account for a significant portion of insoluble aggregates in biochemical assays. In addition, our recent work has implied that [^18^F]PM-PBB3 captures both soluble oligomeric and insoluble fibrillary tau assemblies (*6*).

Besides apoptotic neuronal death evoked by oligomeric and other toxic tau conformers (*55*), living neurons burdened with tau fibrils could undergo phagocytic elimination by aggressive microglia. In our latest observation of living rTg4510 mouse brains with two-photon laser microscopy, live neurons bearing tau deposits were decorated with opsonins such as complements C1q and C3, resulting in engulfment of these cells by activated microglia (*56*). This non-cell-autonomous pathway of neuronal loss initially involves exposure of phosphatidylserine (PS) on the cell surface (*57*), and elevated intracellular calcium levels are known to facilitate the PS externalization (*58, 59*). Depressed susceptibility of excitatory neurons to epileptiform discharges by DSR-143630 would accordingly deactivate opsonization of neurons by phagocytic microglia. In fact, we noted mitigation of reactive astrogliosis and inflammatory microgliosis in rTg4510 mice at 6 months following the treatment over 2 months. Correspondingly, an auxiliary RNAseq analysis has also indicated that expression levels of C1q and C3 were significantly downregulated in the hippocampus of treated animals. The neuroinflammatory response of glial cells did not differ between control and compound-treated groups at 9 – 11 months of age since Iba-1-positive microgliosis in this model plateaued at 8 months (*60*), implying that surviving neurons with densely packed tau filaments do not intensely elicit gliotic responses. Detrimental astroglial and microglial activations might also aggravate the spread of tau pathologies (*61, 62*), although the effects of DSR-143630 on glial modulation of tau dissemination are still enigmatic without evidence from a proper model of tau propagation.

The pharmacological boost of Nav1.1 functions could be advantageous over existing agonists and positive modulators of GABAergic neurotransmissions in the preclusion of neuronal hyperexcitability since constitutive blockade of excitatory signals may give rise to functional deficits of neural networks. The adverse actions of elevated tonic inhibitory activities are exemplified by disturbed attention and cognitions arising from treatments with benzodiazepines and related drugs (*63*). DSR-143630 could repress bursting circuit excitations by fortifying GABA-mediated inhibitory synaptic transmission following the high-frequency repetitive firing of excitatory neurons without altering basal neuronal activities. In principle, conventional anti-epileptic drugs also selectively antagonize epileptogenic prolongation of depolarization due to aberrant ionic trafficking, including glutamate-induced calcium influx and opening of Nav1.6 or Nav1.2 but should barely modify physiological conductions. However, dose-related side effects of these drugs, represented by somnolence, dizziness, and ataxia, commonly occur as a consequence of the general dampening of neuronal activities (*64*). Moreover, target components that are blocked by these drugs could be declined in concurrence with the degeneration of excitatory neurons along with the disease progression, complicating the tuning scheme with minimized adverse events. Although newer anti-epileptic SV2a ligands, levetiracetam and brivaracetam, have shown efficacy for lowering epileptiform activities in AD model mice (*65–67*), it should be noted that SV2a was found to decrease in several regions of the AD brains (*68*), potentially hampering the determination of a drug dosage that is optimal across brain areas. Unlike these agents, there is a plausible rationale for the use of Nav1.1 activators in AD and associated disorders in view of compromised interneuronal integrity.

Our therapeutic paradigm may also be feasible for bursting discharges elicited by a dialog between Aβ and tau in AD (*69*). Indeed, the DSR-143630 administration improved neurocognitive phenotypes in APP transgenic mice (**Fig. S7**), and our pilot experiments indicated a treatment-related diminution of phosphorylated tau accumulation in this model. Notwithstanding these benefits, the selectivity of DSR-143630 for Nav1.1 might not suffice for clinical utilization, and the development of more selective chemical derivatives would offer future biomedical applications with minimal off-target effects. For instance, as Nav1.5 is mainly expressed in the heart, its activation may cause cardiac arrhythmia such as long QT syndrome (*70*). In addition, excitatory neurons predominantly express Nav1.2 and Nav1.6, mediating prolonged electric discharges of these cells. Nevertheless, we found that intraperitoneal administration of 100 mg/kg DSR-143630 significantly suppressed hyperthermia-induced seizures in Dravet mice, supporting the view that DSR-143630 at this dose preferentially activates Nav1.1 rather than other Nav subtypes. The concentration of DSR-143630 in the brain achieved here was also maintained in the treatment of rTg4510 mice (**Table S1 and S2**), validating the dosage for selective stimulation of CNS Nav1.1. In summary, our findings in the tauopathy model indicate that pharmacological activation of Nav1.1 in PV neurons is a promising approach to preventing aberrant brain oscillation, tau fibrillogenesis, and resultant neurodegeneration in AD and allied illnesses. Bimodal tau PET and MRI can accordingly be employed for the longitudinal pursuit of therapeutic efficacies of Nav1.1 activator in living animal models, serving for the selection of potent agents in this class. In conjunction with these monitoring techniques, interneuron-targeting restoration of E/I balances in a disordered brain milieu susceptible to epileptic seizures would offer attractive disease-modifying strategies in clinical settings.

## MATERIALS AND METHODS

### Study design

Mice were housed in a 12 h light/dark cycle with ad libitum access to diet and water. All experimental procedures involving mice were performed with the approval of the Institutional Animal Care and Use Committees in the National Institutes for Quantum and Radiological Sciences and Technology or Sumitomo Pharma Co., Ltd. ddY mice were purchased from SLC Japan. rTg4510 mice were produced by crossbreeding between a parental mutant tau responder line (tetO-MAPT*P301L, FVB/N background) and a tTA activator line (Camk2a-tTA, 129/SV background) as previously described (*27*). In the present study, female rTg4510 mice ranging in age from 4 to 11 months were used for PET and MRI experiments. For the longitudinal assessment of DSR-143630 treatment effects on the tau-associated pathologies, age-matched rTg4510 mice were assigned to the control and treatment groups. DSR-143630 was then administered at 5.8 mg / g-food diet from 4 months until sacrificed for postmortem analysis. Independently, male non-Tg and rTg4510 mice received control or DSR-143630 diet from 4 months of age and were sacrificed for immunohistochemical assessment of reactive gliosis at 6 months of age. A transgenic mouse strain overexpressing human APP751 isoform harboring Swedish and Indiana mutations under the control of Thy1.2 promoter was generated as described previously (*71*). This mouse model typically demonstrates progressive Aβ deposition associated with cognitive impairment. A mouse model for Dravet syndrome (Scn1a(+/−), Balb/c-Scn1a<+/−>) was provided by RIKEN BRC through the National BioResource Project of MEXT, Japan (BRC No.: RBRC06422). This BALB/c strain has a spontaneous mutation in Scn1a, heterozygous mutant mice demonstrate heat-induced epileptic seizures, and homozygous mice die at around 3 weeks old due to severe myoclonic epilepsy in infancy (*72*). We analyzed female heterozygous mutant mice (2-6 months of age) to test the acute effect of DSR-143630 administration.

### Reagents

Nav1.1 activator, DSR-143630, was discovered by a high-throughput screening with Sumitomo Pharma’s compound library and the structure optimization. Mouse monoclonal tau antibodies Tau-5 and Tau-12 were kind gifts from Drs. Lester I. Binder and Nicholas M. Kanaan (Michigan State University). The mouse monoclonal tau antibody PHF1 was kindly provided by Dr. Peter Davies (Albert Einstein College of Medicine). The following antibodies were purchased from commercial sources: mouse anti-tau antibodies Tau46 (A16621), AT8 (MN1020), pS422 (44-764G), and AT180 (MN1040) from Thermo Fisher Scientific; mouse anti-GFAP (MAB360), and guinea pig anti-NeuN (ABN90) from Millipore; mouse anti-βActin (A1978) from Sigma.

### SynchroPatch 768PE

SyncroPatch 768PE (Nanion, München, Germany) was used to generate DSR-143630 potentiation dose responses for Nav channels. Chips with single-hole medium resistance (5−8 MΩ) were used. Pulse generation and data collection were performed with PatchControl384 Version 1.6.6 and DataControl384 Version 1.8.0. Whole-cell recordings were conducted according to Nanion’s standard procedure. Briefly, cells stably expressing human Nav (hNav) channels were stored in a cell hotel reservoir (4°C; shake speed 60 rpm). After initiation of the experiment, cell catching, sealing, whole-cell formation, liquid application, recording, and data acquisition were performed sequentially. The intracellular solution consisted of (in mM) the following: 110 CsF, 10 CsCl, 20 EGTA, and 10 HEPES (pH adjusted to 7.2 with CsOH, and osmolarity adjusted to 285 mOsm); the extracellular solution contained (in mM): 135 NaCl, 4 KCl, 1 MgCl_2_, 5 CaCl_2_, 5 D-glucose monohydrate, 10 HEPES (pH adjusted to 7.4 with NaOH and osmolarity adjusted to 300 mOsm). A holding potential of −100 mV was used; inward currents were recorded using a pulse pattern consisting of a voltage ramp from –100 mV to +85 mV, 35 msec, and a return to −100 mV holding potential. The sweep was repeated every 10 s. The area under the curve (AUC) of each current was analyzed, and the effect of a given compound concentration was calculated as follows: %Potentiation = (IEND/IBASE-1) x100, (IBASE = baseline currents, IEND = currents after compound treatment, an average of six sweeps just before (IBASE) and after (IEND) the compound given). HEK293 cells stably transfected with human Nav1.1 gene were provided by Eurofins DiscoverX Products LLC. (cat#CYL3009). Nav1.4- and Nav1.7 (CT4003)-expressing cells were purchased from ChanTest (currently Charles River Laboratories). Nav1.2- and Nav1.5-stably expressing CHO-K1 cells were established by transfection with a plasmid encoding TET repressor, pcDNA6/TR (Invitrogen), and cDNA of full-length human Nav1.2 or Nav1.5, pcDNA3.1TetO10Zeo. Nav1.6-stably expressing HEK293 cells were established by transfection with a plasmid encoding TET repressor, pcDNA6/TR (Invitrogen), and cDNA of full-length human Nav1.6, pcDNA4/TO.

### Slice preparation and whole-cell patch-clamp recordings

Female Scn1a(+/-) or rTg4510 mice (7-8 weeks old) were decapitated under isoflurane anesthesia, and brains were quickly removed and submerged in an ice-cold cutting solution containing the following (in mM): 87 NaCl, 75 sucrose, 2.5 KCl, 1.25 NaH_2_PO_4_, 0.5 CaCl_2_, 7 MgCl_2_, 25 NaHCO_3_, 25 glucose, 1 Na-pyruvate, and 1 Na-ascorbate, saturated with 95% O_2_/5% CO_2_. Coronal slices (300 μm thick) containing the prefrontal cortex were prepared using a microslicer (VT1000S; Leica). Slices were incubated in a chamber filled with artificial cerebrospinal fluid (ACSF) containing the following (in mM): 125 NaCl, 2.5 KCl, 1.25 NaH_2_PO_4_, 2 CaCl_2_, 1 MgCl_2_, 25 NaHCO_3_, and 25 glucose, 1 Na-pyruvate, and 1 Na-ascorbate, saturated with 95% O_2_/5% CO_2_ at 33–35°C for 30 min, and then placed at room temperature for 30 min before recording.

Slices were placed in a recording chamber on an upright differential interference contrast microscope (Eclipse FN1; Nikon) and continuously superfused with ACSF (room temperature) saturated with 95% O_2_/5% CO_2_. Pipettes were pulled from thin-walled borosilicate glass capillaries with a micropipette puller (Model P-97; Sutter Instrument). Pipette resistance was 4–7 MΩ when pipettes were filled with an internal solution containing the following (in mM): 125 K-gluconate, 10 KCl, 10 HEPES, 10 EGTA, 2 MgCl_2_, 2Na_2_-ATP, 0.3 Na-GTP, and 10 Na_2_-phosphocreatine. The pH value was adjusted to 7.3 with KOH. In the medial prefrontal cortex, neurons were visualized by infrared video microscopy (C3077-79; Hamamatsu Photonics), and non-pyramidal neurons (i.e., putative GABAergic interneurons) were selected for whole-cell patch-clamp recordings. Recordings were performed by Multiclamp 700B amplifier and pCLAMP 10 software (Molecular Devices). In the current-clamp mode, 50 pA-incremental current steps for 400 ms ranging from 0 to 600 pA were injected to record the input-output (I-O) relationship. After recording the I-O relationship in the presence of vehicle solution containing 0.1% DMSO in ACSF, test solution containing DSR-143630 at a concentration of 30 µM in ACSF was superfused for 10 min, and the I-O relationship was recorded again. The number of action potentials evoked by depolarizing current pulses was counted using event-detection with pCLAMP 10 software (Molecular Devices).

### Evaluation of febrile seizure in Scn1a(+/−) mouse

To assess the susceptibility to febrile seizure of Scn1a(+/−) mice, we designed a custom-built observation chamber connected to a water bath heating system in which the body temperature of mice was gradually raised. Mice were placed into the chamber, and the response time (latency) to seizure onset of each mouse was scored. Once we observed seizure onset, the mouse was quickly removed from the chamber, and rectal temperature was measured as an alternative index. For the evaluation of the anticonvulsant effect of DSR-143630 against febrile seizure, the drug was intraperitoneally administered 20 min before placement into the chamber. In case any thermal convulsion was not observed during the 60-min experimental session, the latency score was set as 60 min at maximum.

### Epidural electroencephalography (EEG) recordings

Mice were surgically implanted with stainless steel EEG electrodes under isoflurane anesthesia. Electrodes were positioned over the right primary visual cortex (V1; coordinates: 3.6 mm anterior and 2.3 mm lateral from lambda), posterior parietal cortex (PPC; 2.0 mm anterior, 2.0 mm lateral), and prefrontal cortex (PFC; 2.8 mm anterior, 0.5 mm lateral). An additional electrode was placed over the cerebrum and served as both ground and reference and over the skull above the olfactory bulb. A stainless steel frame (Narishige, Japan) for head fixation was also secured to the skull with dental cement. EEG signals were amplified via a Neuralynx EIB-PTB headstage and recorded using Open Ephys acquisition boards at a sampling rate of either 2 kHz or 30 kHz. Electromyogram (EMG) signals were simultaneously recorded using flexible wire electrodes inserted into the neck muscles. In a subset of animals (3 non-Tg and 3 rTg4510 mice), tetrode recordings were performed by implanting custom-made tetrodes into the right primary visual cortex (V1; AP –3.6 mm, ML +2.3 mm). Tetrode fabrication and implantation procedures followed those described previously (*73*). To assess the acute effects of DSR-143630 on cortical activity, EEG recordings were performed in 4-to 5-month-old rTg4510 and control mice. DSR-143630 (30 mg/kg) or vehicle (0.5% methylcellulose) was administered intraperitoneally 30 minutes before EEG recording during the wheel running task. For chemogenetic suppression of PV neurons, 500 nL of AAV-S5E2-hM4Di-T2A-mScarlet was injected into the same cortical site used for EEG electrode placement (e.g., right V1) via the craniotomy opened during electrode implantation. EEG preprocessing included initial downsampling to 1 kHz with an 8th-order IIR filter, followed by bandpass filtering (1–500 Hz) and removal of line noise. To remove shared noise components, signals were re-referenced by subtracting the average signal across the four recording sites (V1, PPC, PFC, and the skull above the olfactory bulb). Time-aligned EEG segments spanning ±3 s around each event (e.g., reward onset or trial initiation) were extracted for analysis. Spectral power was computed using a multi-taper spectrogram (Chronux toolbox) with a 1-second moving window (step size: 10 ms; time-bandwidth product NW = 1; number of tapers K = 1). Power spectra were log-transformed and averaged across trials. For each animal, spectral power was computed separately for distinct behavioral states (e.g., stop vs. run) based on task-aligned time windows. Baseline power was estimated from pre-stimulus or stopping periods, and movement-related power was normalized against baseline using the decibel scale (10 × log₁₀[movement/baseline]).

### Behavioral task during EEG recordings

During head-fixed EEG recordings, mice were placed on a freely rotating wheel and engaged in a visual-guided operant task under head-fixation. Behavioral control, including stimulus presentation and reward delivery, was implemented using Bpod and Pulse Pal systems (Sanworks, Stony Brook, NY), as described previously (*74*). During head-fixed recording sessions, mice were placed on a freely rotating running wheel under head-fixation, allowing spontaneous transitions between stop and locomotion on a trial-by-trial basis. We defined sessions with more than 20 completed trials as valid sessions and included them in the analysis. On average, control and rTg4510 mice completed 54 ± 25 and 50 ± 25 trials per session, respectively. A display monitor was positioned 15 cm in front of the left eye of the animal. Before trial initiation, the screen remained black. Each trial began when the animal remained stationary on the wheel for 1 second, triggering the presentation of a vertical grating stimulus (alternating black and white stripes on a gray background) on the display as a cue. As the animal rotated the wheel, the grating gradually rotated accordingly, reaching a horizontal orientation after three full wheel turns and then stopping. If the animal remained stationary for another 1 second in this state, a water reward was delivered via a nozzle positioned near the mouth. Overrun was quantified as the ratio of the actual wheel rotation distance during each trial to the threshold distance required to obtain a reward. A ratio of 1.0 indicates that the mouse ran just enough to receive the reward, whereas values greater than 1 reflect excessive running beyond the required threshold.

To assess the acute effect of PV interneuron suppression on cortical activity, DCZ (0.1 mg/kg, i.p.) was administered 5 minutes before recording. For pharmacological intervention, DSR-143630 (30 μM) was freshly prepared and administered intraperitoneally 30 minutes before EEG recording.

### Recombinant adeno-associated virus for chemogenetic suppression of PV neurons

pAAV-S5E2-GFP-fGFP was a gift from Jordane Dimidschstein (Addgene plasmid #135631) pAAV-GFAP-hM4D(Gi)-mCherry and pAAV-hSyn-mScarlet were kind gifts from Bryan Roth (Addgene plasmid # 50479) and Karl Deisseroth (Addgene plasmid # 131001), respectively For constructing a plasmid of pAAV-S5E2-hM4Di-T2A-mScarlet, cDNAs encoding each protein were individually amplified by PCR and reassembled by replacing GFP-fGFP sequence of pAAV-S5E2-GFP-fGFP. The sequence of the plasmid DNA was verified before being used for transfection. Recombinant AAVs were produced and packaged with serotype DJ in HEK293T cell cultures and purified by AAVpro® Purification Kit (TaKaRa). The virus titer was determined by AAVpro® Titration kit (for Real Time PCR) ver.2 (TaKaRa) as described previously (*25*). For validation of AAVs targeting PV neurons, C57BL/6j mice at 2 months of age were anesthetized with isoflurane and received stereotaxic injection of 1.0 μL AAVs (at a titer ranging from 3.4 x10^11^ to 1.4 x10^12^ vg/ml) into a side of the somatosensory cortex for histological analysis. To assess the functional impact of PV neuron suppression, control littermates of rTg4510 mice (lacking both tetO-MAPT*P301L and Camk2a-tTA transgenes) at 12 weeks of age received 1.0 μL injections of AAV-S5E2-hM4Di-T2A-mScarlet (1.4 × 10¹¹ to 3.2 × 10¹¹ vg/mL) into either the primary visual cortex (V1; AP –3.6 mm, ML +3.2 mm) or posterior parietal cortex (PPC; AP –2.0 mm, ML +2.0 mm). A stainless-steel screw EEG electrode was simultaneously implanted at the same coordinates as the injection site. After a 4-week expression period, mice were tested in a head-fixed wheel-running task. Saline or DCZ (1.0 mg/kg, intraperitoneal) was administered 5 minutes before recording, and EEG signals were collected during behavioral performance.

### MRI

All MRI experiments were performed on a 7.0 T horizontal MRI scanner (Magnet: Kobelco and JASTEC, Japan; Console: Bruker Biospin, Germany) with a volume coil for transmission (Bruker Biospin) and a quadrature surface coil for reception (Rapid Biomedical, Germany). The mice were initially anesthetized with 3.0% isoflurane (Escain, Mylan Japan, Japan), and they were maintained with 1.5 ∼ 2.0% isoflurane and 1:5 oxygen/ room-air mixture during the MRI experiments. Rectal temperature was continuously monitored by an optical fiber thermometer (FOT-M, FISO, Canada) and maintained at 36.5 ± 0.5°C using a heating pad (Rapid Biomedical) and warm air. The first imaging slices were carefully set at the rhinal fissure based on the mouse brain atlas. Trans-axial T2-weighted fast spin-echo MR images were acquired using a rapid acquisition with relaxation enhancement (RARE) sequence. The imaging parameters were as follows: repetition time (TR) = 4,200, echo time (TE) = 36 ms, Fat-suppression = on, number of averages (NA) = 4, RARE factor = 8, number of slices = 28, FOV = 25.6 mm × 14.5 mm, matrix = 256 × 256 and scan time = 14 min. Frequency-selective saturation pulses and crusher magnetic field gradients were used for fat suppression.

### PET imaging

Radiosynthesis of a tau PET tracer, [^18^F]PM-PBB3, was performed as described previously (*6*). Before PET imaging, rTg4510 mice were deeply anesthetized with 1.5 - 2.0% isoflurane, and a 30-gauge needle with a catheter was inserted into the tail vein for injection of [^18^F]PM-PBB3. Mice were placed on the stage of a microPET Focus 220 animal scanner (Siemens Medical Solutions, Malvern, PA) providing 95 transaxial planes 0.815 mm (center-to-center) apart, a 9.0-cm transaxial field-of-view (FOV), and a 7.6-cm axial FOV. Immediately after a bolus administration of [^18^F]PM-PBB3, an emission scan in 3D list mode was conducted for 60 min. All list-mode data were sorted into 3D sinograms and were then Fourier-rebinned into 2D sinograms ([^18^F]PM-PBB3; 1 min x 4, 2 min x 8, 5 min x 8). Images were reconstructed with filtered back-projection using a Hanning filter with a Nyquist cutoff of 0.5 cycle/pixel.

### MRI and PET data analysis

For measurement of regional brain volumes, the anatomical volume of interest (VOI) was manually segmented on the T2-weighted MRI in each slice to define the neocortex, hippocampus, striatum, cerebellum, and lateral ventricle using PMOD® software (ver. 3.8, PMOD Technologies Ltd., Zurich, Switzerland). Brain contours were double-checked by another investigator to finalize the registration of VOIs. For quantification of regional radioactive signals of [^18^F]PM-PBB3, dynamic PET images were co-registered with MR images with defined anatomical VOIs using PMOD software. The tracer uptake in each VOI was estimated as standardized uptake value [SUV = radioactivity concentration (kBq/ccm) / injected dose (kBq) compensated by body weight (g)]. The specific binding of the tracer in each VOI was analyzed by calculating the SUV ratio (SUVR) to the reference region (cerebellum).

### Histological analysis

9-11-month-old female rTg4510 mice were sacrificed by cervical dislocation, and the brain tissues were immediately removed and fixed in 4% paraformaldehyde followed by cryoprotection in 20% sucrose in phosphate buffer saline. Brain tissues were then embedded with OTC compound, and 20-μm-thick frozen sagittal slices were cut on an HM560 cryotome and attached to a glass slide. For immunohistochemical analysis, slices were autoclaved for antigen retrieval, blocked, and incubated with primary antibodies against NeuN and pTau AT8. Alternatively, 6-month-old male rTg4510 mice were deeply anesthetized and sacrificed by 4% PFA perfusion. Brains were dissected, fixed, and cut to 50-μm free-floating coronal slices with a Leica VT1200S vibratome. All slices were blocked in PBS-supplemented 4% BSA, 2% horse serum, and 0.25% Triton X-100 at room temperature and then immunostained with anti-GFAP antibody. After probing with Alexa Fluor 488 or 546 conjugated secondary antibodies, the slices were mounted in VECTASHIELD medium (Vector Laboratories). Fluorescence images were captured using Keyence BZ-X710 fluorescence microscopy (Keyence) with either x10 (Plan-Apo, NA 0.45) or x20 (Plan-Apo, NA 0.75) objectives or by LSM880 laser scanning confocal microscopy (Carl Zeiss) with either x10 (Plan-Apo, NA 0.45) or x20 (Plan-Apo, NA 0.8) objectives. For identification of neurofibrillary tangle-like tau deposition, slices were analyzed with Gallyas-Braak silver staining after pretreatment with 0.25% KMnO_4_ followed by 2% oxalic acid as described previously (*77*).

All quantification analysis of acquired images was performed using Image J software. For assessment of the cortical area and neuronal density of brain slices, a wide range of tiled images labeled with anti-NeuN antibody was first generated, and regions of interest (ROIs) were manually set on the neocortical region for determining the total cortical area. High magnification images were next captured in the region of the sensory-motor cortex and converted to binary images by thresholding, and the number of NeuN^+^ cells was automatically calculated by Analyze Particles function. To minimize the variability between the different levels of each slice, data from 2-4 slices from each brain were averaged for this analysis. For quantification of immunoreactive signals of anti-pTau AT8, and anti-GFAP antibodies in the sensory cortex and hippocampal CA1, each brain region on the captured image was manually segmented by ROI analysis. The average intensity/pixel was then measured to determine the total immunoreactivity of each of the antibodies.

### Biochemistry

Biochemical extraction and fractionation of brain tissues were performed as described previously (*29*). Briefly, a frozen hemisphere of brain tissue was homogenized in 10 volumes of Tris-buffered saline supplemented with protease and phosphatase inhibitors (Sigma, St. Louis, MO) and centrifuged at 27,000 g for 20 min to obtain the supernatant (S1) and pellet. The pellets were then re-suspended in 5 volumes of high salt/sucrose buffer and further centrifuged at 27,000 g for 20 min. The resultant supernatant was then extracted in 1% sarkosyl for 60 min, followed by ultra-centrifugation at 150,000 g to the supernatant (S3) and pellet (P3) fractions. Sarkosyl-insoluble proteins in P3 pellets were recovered in SDS-sample buffer and stored at -20°C until use. For assessment of tau solubility by western blot, proteins in S1 and P3 were separated by SDS-PAGE, transferred onto nitrocellulose membrane, probed with primary and secondary antibodies, and analyzed by Amersham Imager 600 (GE Healthcare) following previously described methods (*29*).

### Novel object recognition test

The novel object recognition test was conducted for the assessment of episodic memory. On the day of the experiment, mice were transferred to the testing room with the same environmental conditions as their housing room. The novel object recognition test consisted of the first trial (T1) and the second trial (T2). The two trials were separated by an inter-trial interval of 3 hr. In T1, each mouse was put in the test box, in which two identical objects (familiar objects) were placed in two adjacent corners, and the time spent exploring each object was measured for 5 min. In T2, the one object explored by each mouse for a shorter period in T1 was replaced by a novel object, and the time spent exploring the familiar object and the novel object was recorded for 5 min. Exploration of an object was defined as sniffing and/or touching with the nose. For the evaluation of acute pro-cognitive effect, DSR-143630 was injected intraperitoneally into APP-Tg mice 30 min before T1. For evaluation of chronic effect, DSR-143630 at 0.75 mg/g-food and 1-Aminobenzotriazole (ABT; an inhibitor of cytochrome P450) at 1 mg/g-food were co-administered to 4-month-old rTg4510 mice for 2 months. Administration of DSR-143630 at 0.75 mg/g-food with ABT achieved equivalent exposure to DSR-143630 administration at 5.8 mg/g-food alone.

### Statistical analyses

Statistical analysis was performed with SAS9.4 (SAS Institute Inc.), MATLAB (The MathWorks, Inc.), or GraphPad Prism (GraphPad Inc., CA) software. For comparison of two rTg4510 groups treated with control and DSR-143630, or comparison between non-Tg group and rTg4510 group, Student t-test was used. For multiple comparisons of various experimental conditions, one-way ANOVA followed by Dunnett post hoc was used to determine differences between the control and other groups. To examine the protective effect of DSR-143630 treatment on tau pathology and brain atrophy, linear regression analysis with ANCOVA was used to determine significant differences. Regarding the effects of DSR administration on behavior and EEG (Fig. 2E, F), because the number of valid sessions obtained differed across animals, we used a linear mixed-effects model with animal as a random effect to account for within-subject correlation and unbalanced sampling, and tested fixed effects (Genotype, Injection, and their interaction). Null hypotheses were rejected at the 0.05 level of *P* values.

## List of Supplementary Materials

Fig S1 to S7

Tables S1 to S2

## Acknowledgments

The authors thank Dr. Ichio Aoki, Takeharu Minamihisamatsu, and Shoko Uchida for technical assistance. We are also grateful to Drs. Yoshio Sakurai and Shuzo Sakata for their valuable advice on electrophysiological data.

## Author contributions

Conceptualization: KS, JH, HT, MH, MS

Investigation: KS, JH, TK, CS, HT, JM, MO, MT, NN, SH, TK, YM, TI, YT

Funding acquisition: JH, MS, MH

Supervision: MRZ, TM, NS, MH

Writing – original draft: KS, JH, TK, HT, NS, MH, MS

## Data and materials availability

All data are available in the main text or the supplementary materials.

