## Supplementary materials for "Pharmacological potentiation of Nav1.1 channels in interneurons mitigates tau depositions and neuronal death in a mouse model of neurodegenerative dementias"

Kazuaki Sampei *et al*.

**The PDF file includes:**

Figs. S1 to S7
Tables S1 to S2

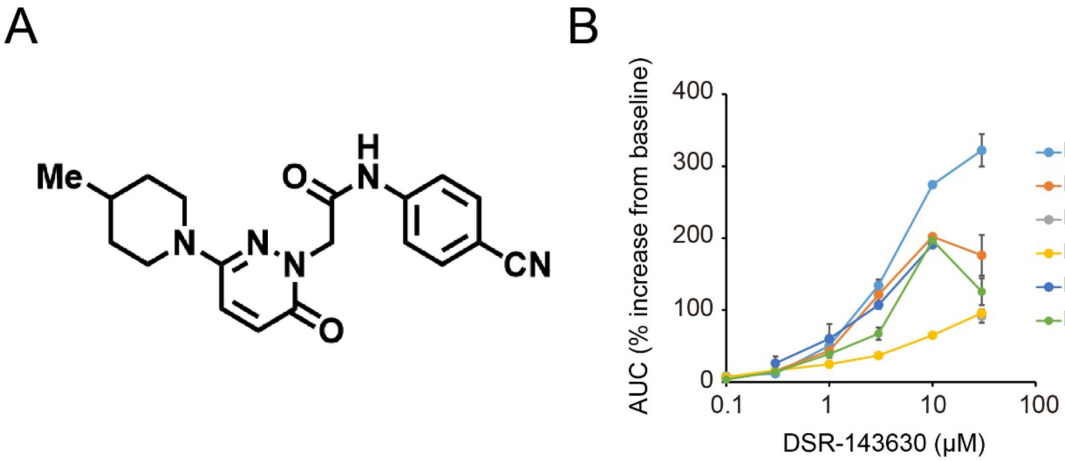


**Fig. S1. The structure and pharmacological effect of DSR-143630**

(A) The chemical structure of DSR-143630.

(B) Concentration-response curves of DSR-143630 on six hNav subtypes. EC50 values for Nav1.1, Nav1.2, Nav1.5, Nav1.6, and Nav1.7 were 3.39 μM, 2.42 μM, 4.87 μM, 1.70 μM, and 1.46 μM, respectively. DSR-143630 potentiated Nav1.1 most strongly at 30 μM. Data are presented as Mean ± S.E.M.

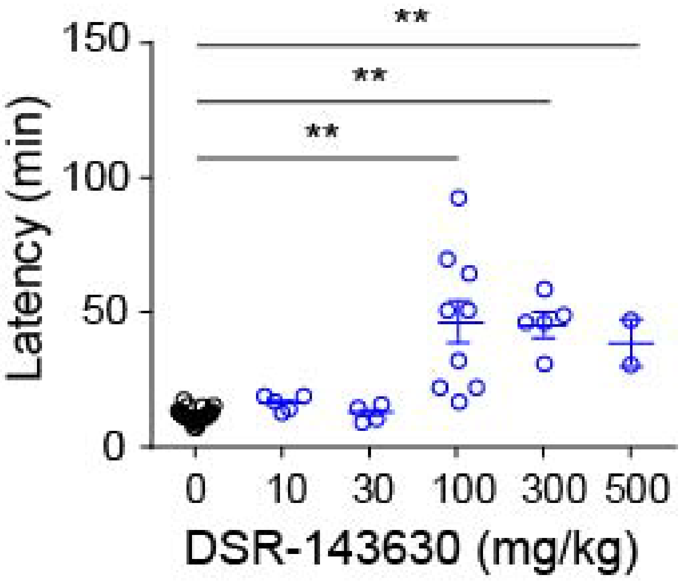


**Fig. S2. Latency to heat-inducible seizure onset of mice receiving various concentrations of DSR-143630**

Scn1a(+/−) mice received various concentrations of DSR-143630 by intraperitoneal injection and were placed into a heating chamber to assess seizure behavior. The graph shows the latency to heat-induced seizure onset of mice receiving different concentrations of DSR-143630. A 0.5% MC solution was administered intraperitoneally as a vehicle control. Data from indicated numbers of animals in each condition are shown as Mean ± S.E.M. **$p < 0.01$ vs. vehicle; parametric Dunnett's multiple-comparison test.

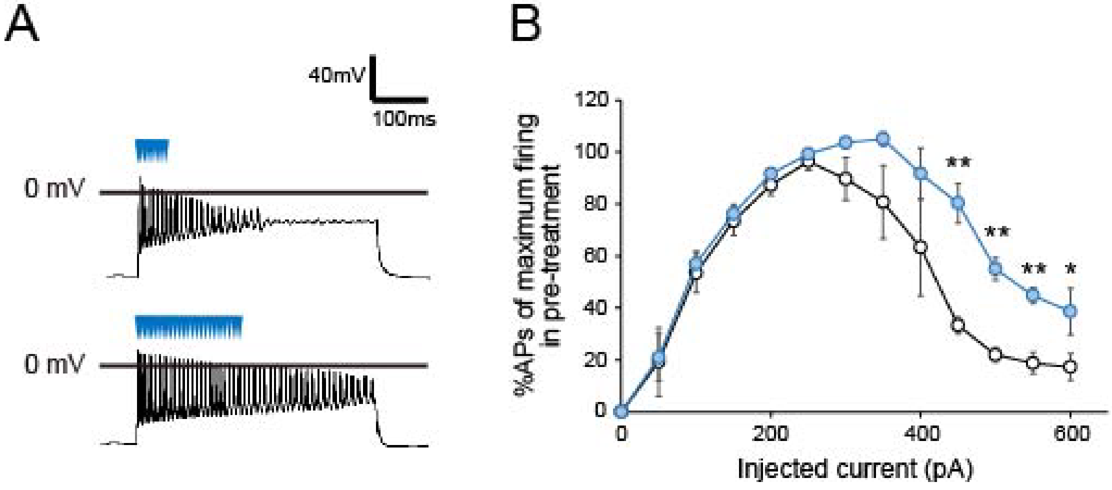


**Fig. S3. Pharmacological effect of DSR-143630 on electrophysiological properties of cortical slices derived from rTg4510 mice**

**(A)** Representative recordings of rTg4510 brain slices with current injection at 600 pA. **(B)** Injected current–response curves of action potential frequencies during 400 ms. The graph shows the percentage of action potentials observed under each experimental condition relative to the maximum number of action potentials observed during pretreatment with 0.1% DMSO in each slice. Curves in black and blue indicate 0.1% DMSO and 30 μM DSR-143630 treatment, respectively. Data are presented as Mean ± S.E.M. (n = 3).

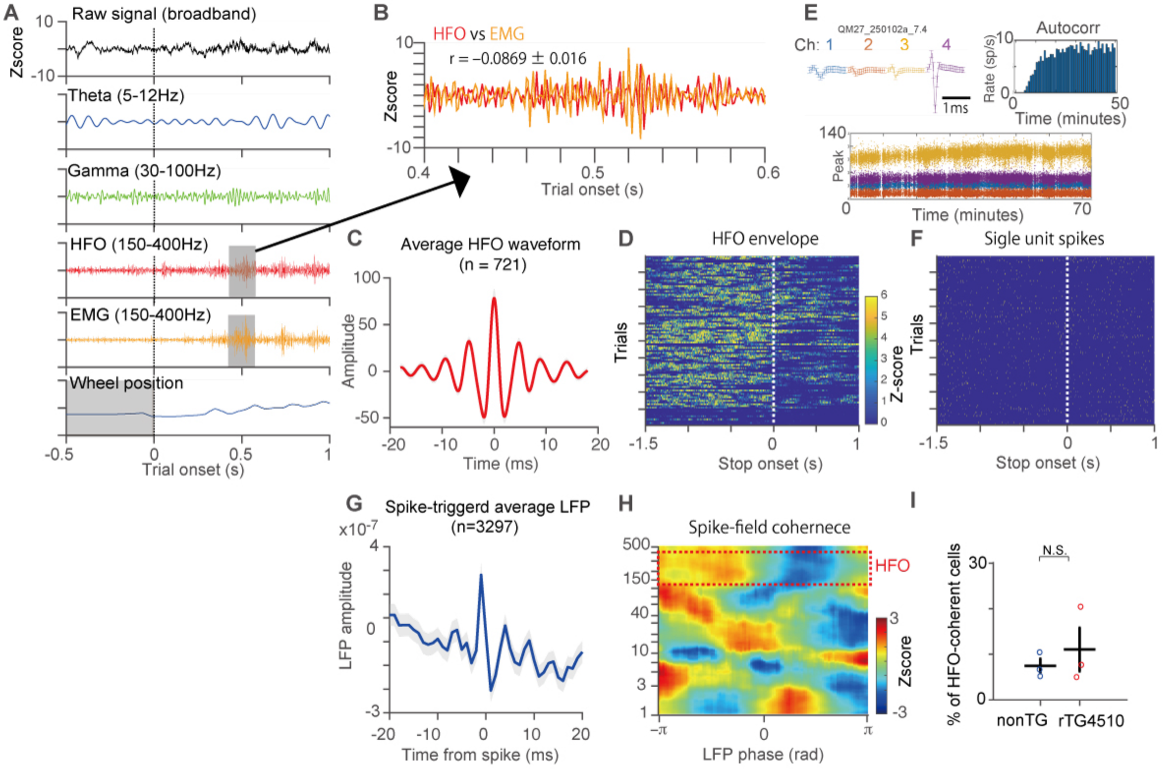


**Fig. S4. Validation of HFOs reflecting intrinsic cortical activity**

**(A)** Broadband LFP signal (top) and bandpass-filtered traces (theta: 5–12 Hz; gamma: 30–100 Hz; high-frequency oscillations [HFO]: 150–400 Hz) aligned to trial onset (time = 0) in an example session. EMG (150–400 Hz) trace and wheel position are also shown. **(B)** Overlaid HFO and EMG traces during a representative trial demonstrate low temporal correlation (mean r = –0.09 ± 0.02). **(C)** The average waveform of detected HFO events (n = 721) reveals multi-cycle oscillations with a typical waveform structure. **(D)** HFO envelope aligned to wheel-stop onset across trials shows trial-by-trial consistency of state-dependent HFO modulation (Z-scored per session). **(E)** An example of a well-isolated single unit from V1 of rTg4510 mouse. **(F)** Single-unit spike raster relative to wheel-stop onset. **(G)** Spike-triggered average of the HFO-filtered LFP (150–400 Hz), aligned to spike times (n = 3,297), shows a structured multi-cycle LFP response. **(H)** Spike-field coherence map (spike phase vs. frequency) reveals phase-locked activity in the HFO range. **(I)** Percentage of single units exhibiting significant HFO-phase locking (Z > threshold) in nonTG and rTG mice (n = 3 animals per group).

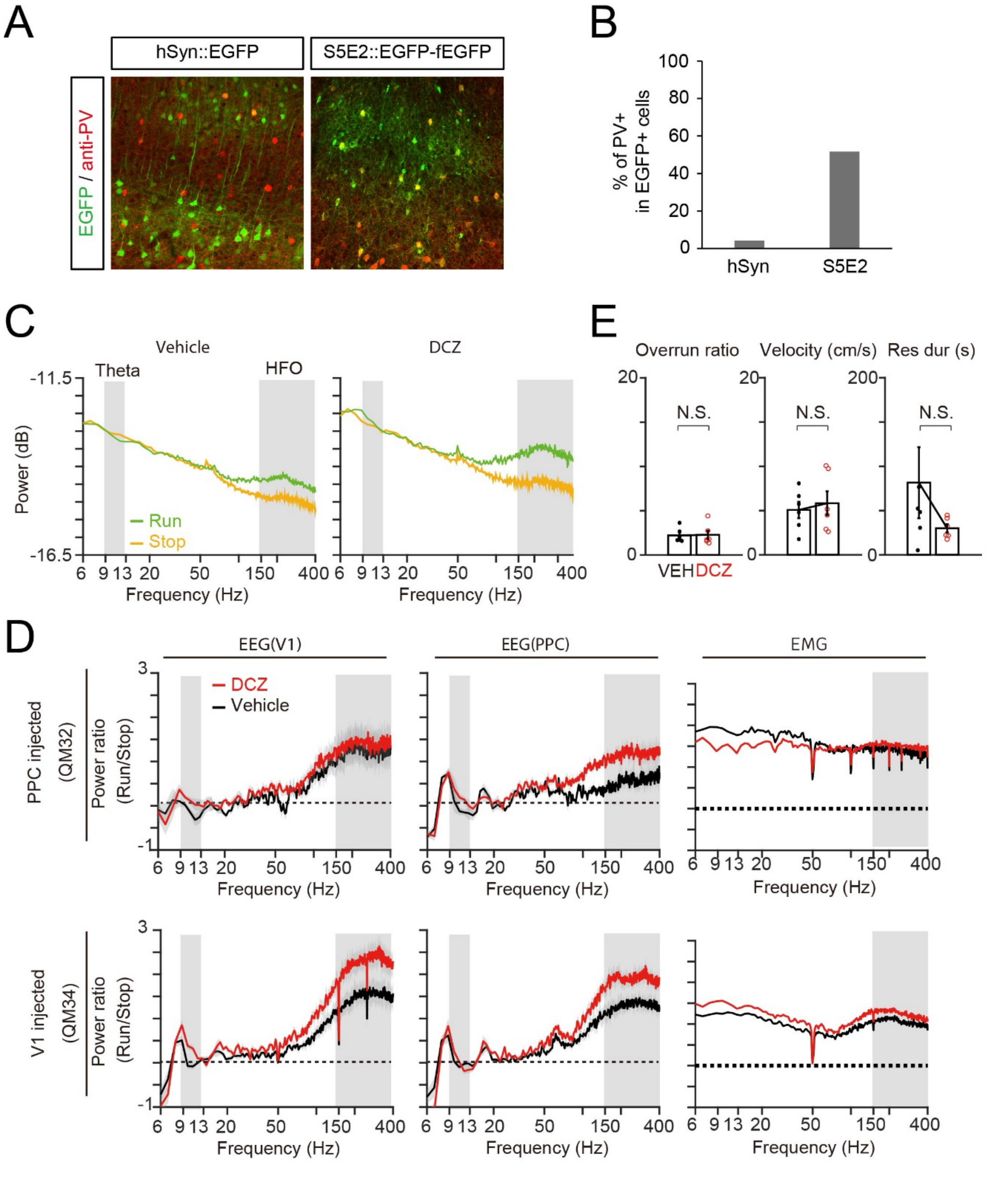


**Fig. S5. Chemogenetic inactivation of cortical PV neurons disturbs brain oscillation and increases HFOs**

**(A)** Immunohistochemical validation of neuronal subtype-specific expression of EGFP under control of S5E2 enhancer. Representative images of EGFP (green) and PV neurons (red) in the somatosensory cortex captured by confocal microscopy are shown. **(B)** Percentage of $EGFP^+$ cells co-labeled with anti-NeuN and anti-PV immunoreactivity. Data from N mice are presented as Mean ± S.D. **p < 0.01 (Student t-test). **(C)** Example EEG power spectral densities during running (green) and stopping (orange) following vehicle (left) or DCZ (1 mg/kg, right) injection. Shaded regions indicate theta (6–13 Hz) and HFO (150–400 Hz) bands. **(D)** Run/stop power ratio spectra recorded from V1 (left), PPC (middle), and EMG (right) in mice injected with AAV-hM4Di-DREADD into PPC (QM32, top) or V1 (QM34, bottom). DCZ (red) increased HFO power in the injected region compared to the vehicle (black); no significant increase was observed in EMG. **(E)** Behavioral parameters (overrun ratio, velocity, response duration) were not significantly affected by DCZ.

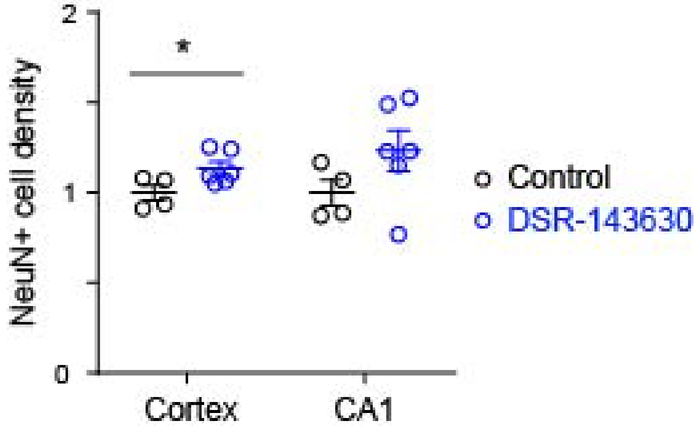


**Fig. S6. Neuronal density in cortex and hippocampus of rTg4510 brains treated with control diet and DSR-143630 diet**

Densities of $NeuN^+$ neurons in hippocampus CA1 and cortex of 9-11-month-old rTg4510 mice chronically administered with control diet or DSR-143630 diet were analyzed by immunohistochemistry. Data from control group (black, n = 4) and DSR-143630 group (blue, n = 6) are presented as Mean ± S.E.M. Each circle represents an individual mouse. *$p < 0.05$, (Student t-test).

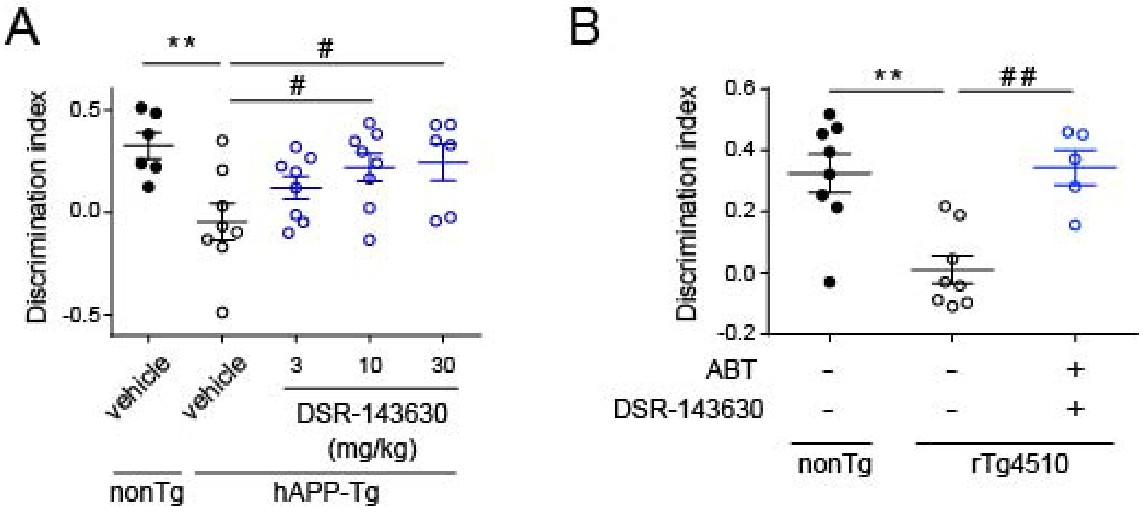


**Fig. S7. Administration of DSR-143630 rescued the cognitive impairment of human APP transgenic model mice and rTg4510 model mice.**

**(A)** The cognitive function of human APP transgenic mice receiving acute administration of DSR-143630 was analyzed by the novel object recognition test. Data from nonTg group (n = 6) and hAPP Tg group received i.p. injection of 0mg/kg (n = 8), 3mg/kg (n = 8), 10mg/kg (n = 8), or 30mg/kg (n = 8) DSR-143630 are presented as Mean ± S.E.M. Each circle represents an individual mouse. **p < 0.01, vs nonTg (Student t-test). #p < 0.05, vs vehicle control (Dunnett's test)

**(B)** Novel objective recognition test of 6-month-old rTg4510 mice chronically treated with DSR-143630 for 2 months. Data from non-Tg group (n = 8) and rTg4510 group with (n = 8) or without (n = 5) DSR-143630 are presented as Mean ± S.E.M. **p < 0.01 (Student t-test). ##p < 0.01, vs vehicle (Dunnett's test)

Table S1

| Animal # | Plasma conc. (μM) | Brain concentration of protein bound fraction (μM) | Brain concentration of unbound fraction (μM) | Kp $_{brain}$ |
|---|---|---|---|---|
| #1 | 37.7 | 18.7 | 0.673 | 0.50 |
| #2 | 34.5 | 16.2 | 0.583 | 0.47 |
| #3 | 39.8 | 21.6 | 0.779 | 0.54 |
| mean | 37.4 | 18.8 | 0.70 | 0.50 |
| SD | 2.66 | 2.73 | 0.10 | 0.04 |

**Table S1. Plasma and brain concentration of DSR-143630 in ddY mice receiving intraperitoneal administration of the drug**

ddY mice were sacrificed 60 min after intraperitoneal injection of 100mg/kg DSR-143630. Plasma and brain tissues were quickly removed, and the concentration of the drug in each fraction was determined by MS-based analysis.

Table S2

| Animal # | Plasma conc. (μM) | Brain concentration of protein bound fraction (μM) | Brain concentration of unbound fraction (μM) | Kp brain |
|---|---|---|---|---|
| #1 | 52.0 | 40.3 | 0.972 | 0.78 |
| #2 | 14.3 | 10.2 | 0.246 | 0.71 |
| #3 | N.C.# | 34.9 | 0.842 | N.C.# |
| #4 | 54.8 | 46.0 | 1.110 | 0.84 |
| #5 | 37.8 | 25.7 | 0.619 | 0.68 |
| #6 | 26.8 | 20.8 | 0.500 | 0.78 |
| mean | 37.1 | 29.7 | 0.71 | 0.76 |
| SD | 17.0 | 13.3 | 0.32 | 0.06 |

**Table S2. Plasma and brain concentrations of DSR-143630 in mice chronically treated with the drug for 4 months and used in MR and tau imaging**

Plasma and brain tissues were quickly isolated from rTg4510 mice chronically treated with DSR-143630 at 9-11 months of age, and the concentration of the drug in each fraction was determined by MS-based analysis. Brain concentration of the unbound fraction of the compound was calculated using the protein binding ratio (97.59%).

#: Because animal #3 died after the final PET imaging, the plasma sample was unavailable. A brain sample from animal #3 was obtained postmortem. N.C.: not calculated.
